# Genes influencing phage host range in *Staphylococcus aureus* on a species-wide scale

**DOI:** 10.1101/2020.07.24.218685

**Authors:** Abraham G. Moller, Kyle Winston, Shiyu Ji, Junting Wang, Michelle N. Hargita Davis, Claudia R. Solis-Lemus, Timothy D. Read

## Abstract

*Staphylococcus aureus* is a human pathogen that causes serious diseases ranging from skin infections to septic shock. Bacteriophages (“phages”) are both natural killers of *S. aureus*, offering therapeutic possibilities, as well as important vectors of horizontal gene transfer in the species. Here, we used high-throughput approaches to understand the genetic basis of strain-to-strain variation in sensitivity to phages, which defines the host range. We screened 259 diverse *S. aureus* strains covering more than 40 sequence types for sensitivity to eight phages, which were representatives of the three phage classes that infect the species. The phages were variable in host range, each infecting between 73 and 257 strains. Using genome-wide association approaches, we identified putative loci that affect host range and validated their function using USA300 transposon knockouts. In addition to rediscovering known host range determinants, we found several previously unreported genes affecting bacterial growth during phage infection, including *trpA, phoR, isdB, sodM, fmtC*, and *relA.* We used the data from our host range matrix to develop predictive models that achieved between 40 and 95% accuracy. This work illustrates the complexity of the genetic basis for phage susceptibility in *S. aureus* but also shows that with more data, we may be able to understand much of the variation. With a knowledge of host range determination, we can rationally design phage therapy cocktails that target the broadest host range of *S. aureus* strains and address basic questions regarding phage-host interactions, such as the impact of phage on *S. aureus* evolution.

**Importance:** *Staphylococcus aureus* is a widespread, hospital- and community-acquired pathogen, many strains of which are antibiotic resistant. It causes diverse diseases ranging from local to systemic infection and affects both the skin and many internal organs, including the heart, lungs, bones, and brain. Its ubiquity, antibiotic resistance, and disease burden make new therapies urgent. One alternative therapy to antibiotics is phage therapy, in which viruses specific to infecting bacteria clear infection. In this work, we identified and validated *S. aureus* genes that influence phage host range - the number of strains a phage can infect and kill - by testing strains representative of the diversity of the *S. aureus* species for phage host range and associating strain’s genome sequences with host range. These findings together improved our understanding of how phage therapy works in the bacterium and improve prediction of phage therapy efficacy based on the infecting strain’s predicted host range.

## Introduction

There is no licensed vaccine for *Staphylococcus aureus* and many clinical strains are resistant to multiple antibiotics. For these reasons, alternative treatments such as bacteriophage therapy are being actively investigated (1, 2). Phage therapy has some advantages over using antibiotics. Phages show little or no human toxicity and the high diversity of natural phages available to be isolated for treatment suggests that complete resistance would be hard to evolve (3, 4). However, there is no natural phage known to kill all *S. aureus* strains and for that reason phage cocktails (mixtures of phages with non-overlapping host ranges) are necessary. Rational cocktail formulation requires comprehensive knowledge of the genetic factors that influence phage host range.

*S. aureus* phages and corresponding known host mechanisms regulating phage resistance and host range have previously been reviewed (1, 5, 6). Known *S. aureus* phages belong to the order *Caudovirales* (tailed phages) and are further divided into three morphological classes - the long, noncontractile-tailed *Siphoviridae*, the long, contractile-tailed *Myoviridae*, and the short, noncontractile-tailed *Podoviridae* (5). The *Siphoviridae* are temperate, while the *Myo*- and *Podoviridae* are virulent (5). The *Siphoviridae* bind either α-O-GlcNAc or β-O-GlcNAc attached at the 4 positions of wall teichoic acid (WTA) ribitol phosphate monomers, while the *Podoviridae* bind only β-O-GlcNAc-decorated WTA, and the *Myoviridae* bind either the WTA ribitol-phosphate backbone or β-O-GlcNAc-decorated WTA (1, 7, 8). *S. aureus* is known to produce poly-ribitol-phosphate rather than poly-glycerol-phosphate WTA (9). WTA biosynthesis genes are conserved throughout the species, with the exception of the unusual sequence type ST395 (10), as are WTA glycosyltransferases *tarM* and *tarS*, but occasional *tarM* inactivation or absence provides *Podoviridae* susceptibility (11).

Currently identified resistance mechanisms in *Staphylococcus* species act at the adsorption, biosynthesis, and assembly stages of infection (1). Adsorption resistance mechanisms include receptor alteration, removal, or occlusion by large surface proteins or polysaccharides (capsule) (7, 11–16). Biosynthesis resistance mechanisms include halting the infection process through metabolic arrest (abortive infection) and adaptive (CRISPR) or innate (restriction-modification) immunity to phage infection through phage DNA degradation (17–21). Temperate phage and staphylococcal pathogenicity islands (SaPIs) inserted in the genome may also offer barriers to *Siphoviridae*, through superinfection immunity and assembly interference, which occurs through SaPI parasitization of the infecting viruses packaging machinery (22–27).

While previous studies have identified numerous individual host resistance mechanisms in *S. aureus*, few have examined the importance of these mechanisms on a species-wide scale. In addition, although many *S. aureus* phages are reported to have wide host ranges (28–34), and even early studies suggested staphylococcal phage therapies to be highly effective (35), experiments conducted thus far have failed to explain the genetic bases of host range or resistance development in a species-wide manner. Only one previous study has associated genetic factors (gene families) with phage resistance using a hypothesis-free method (36). This work used a two-step linear regression model to associate some 167 gene families, mostly of unknown function, with resistance assessed in 207 clinical MRSA strains and 12 phage preparations. However, the study did not associate any other types of genetic changes with host range and examined only MRSA strains.

In this study, we associated multiple genetic factors - gene presence/absence, point mutations, and more complex polymorphisms - with *S. aureus* phage host range and resistance in a hypothesis-free, species-wide, and genome-wide manner. We used a novel high-throughput assay to determine resistance phenotypes of 259 strains challenged with eight *S. aureus* phages belonging to all three morphological categories (*Siphoviridae*, *Myoviridae*, and *Podoviridae*). We then used two bacterial genome-wide association study techniques to identify core genome single nucleotide polymorphisms (SNPs), and subsequences of length k (k-mers) significantly associated with each phenotype and used these significant features to develop predictive models for each phenotype. We also tested for associations between phenotypes and phylogeny, clonal complex (CC), and methicillin resistance (MRSA) and validated novel genes found to be associated with sensitivity or resistance in the GWAS through molecular genetics, thus complementing the hypothesis-free GWAS approach with hypothesis-driven experiments and demonstrating that GWAS-discovered determinants have causative effects on phage resistance.

## Materials and Methods

### Strains, media, and phage propagation

Phages used in this study were phage p0045 (80α-like), p0017S, p002y-DI p003p-Mourad 87, and p0040-Mourad 2 (*Siphoviridae*); p0006-K and pyo (*Myoviridae*); and p0017-HER49/p66 (*Podoviridae*). All phage genomic DNA was isolated with the bioWORLD Phage DNA Isolation Kit following manufacturer’s directions after phage precipitation by a previously described protocol. The corresponding genomes were prepared for sequencing with a 1D ligation sequencing kit (SQK-LSK109) or 1D rapid sequencing kit (SQK-RAD004) and sequenced with an Oxford Nanopore MinION using a Flongle flow cell (FLO-FLG001). Phage p0045, p0017S, p002y-DI, p003p-Mourad 87, p0040-Mourad 2, and p0006-K genomes were also sequenced with Illumina technology by the Microbial Genome Sequencing Center (MiGS) at the University of Pittsburgh.

All *Siphoviridae* and *Myoviridae* were propagated in *S. aureus* RN4220, while the sole podovirus was propagated in *S. aureus* RN4220 *tarM*::Tn, which was constructed by transducing strain RN4220 with Nebraska Transposon Mutant Library (NTML) (37) strain USA300 JE2 *tarM*::Tn (NE611) phage 0045 lysate. Strains, phages, and plasmids used for phage propagation and molecular genetic validation of GWAS results are listed in **Table 1**. Transduction was performed according to a previously published protocol (38). All overnight cultures were grown in LB/TSB 2:1 supplemented with 5 mM CaCl_2_ to promote phage adsorption.

**Table 1:**
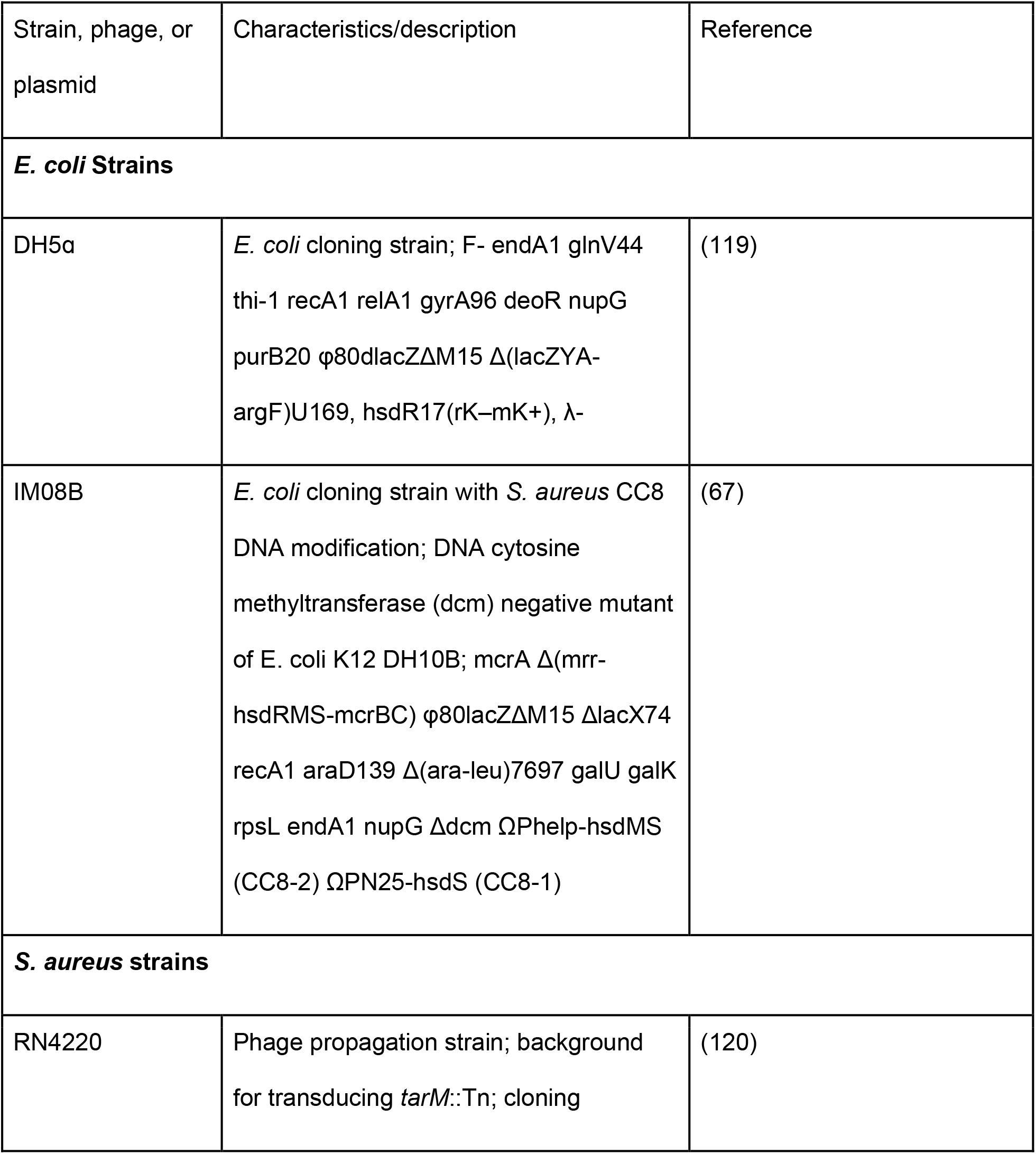
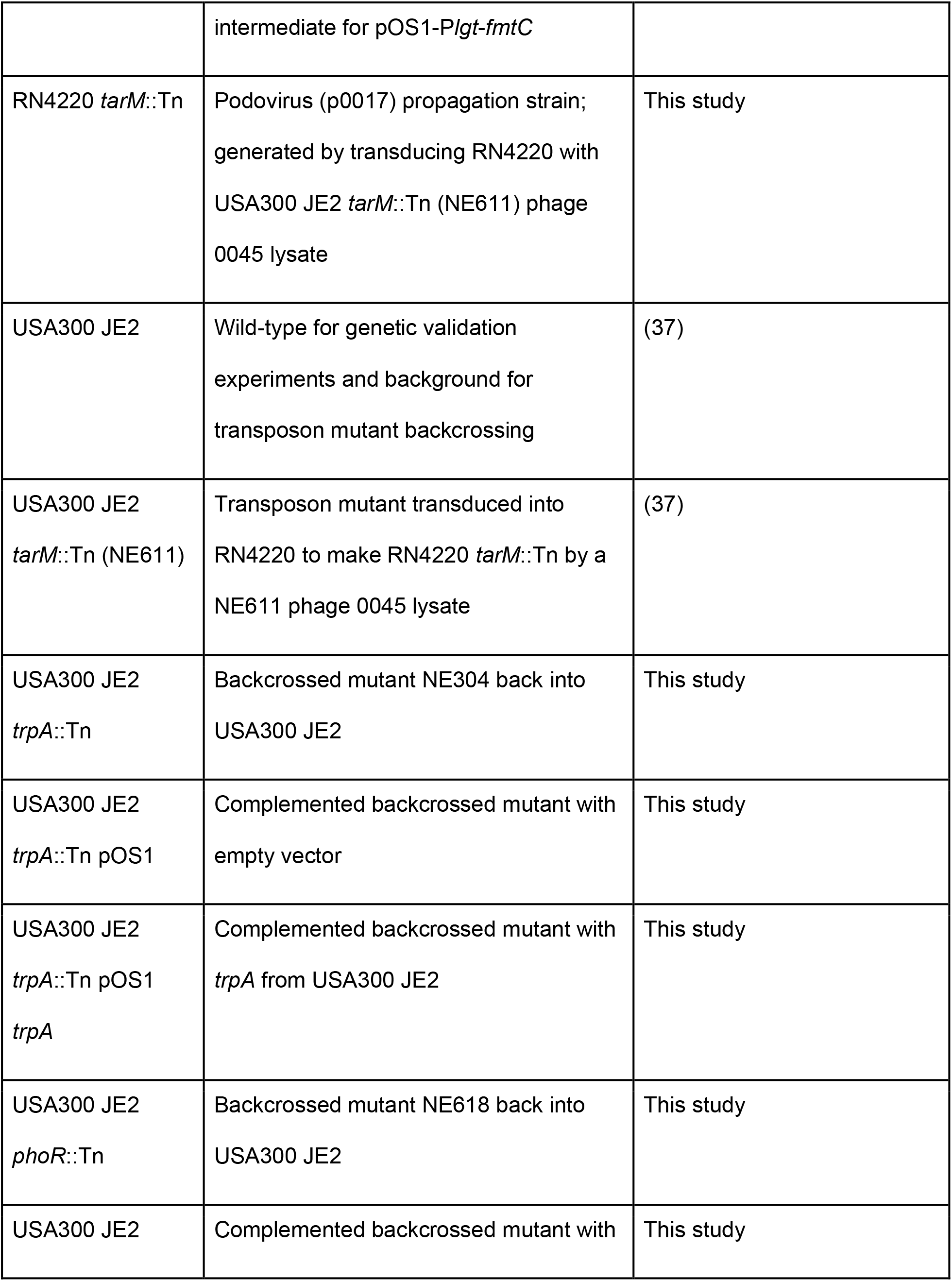
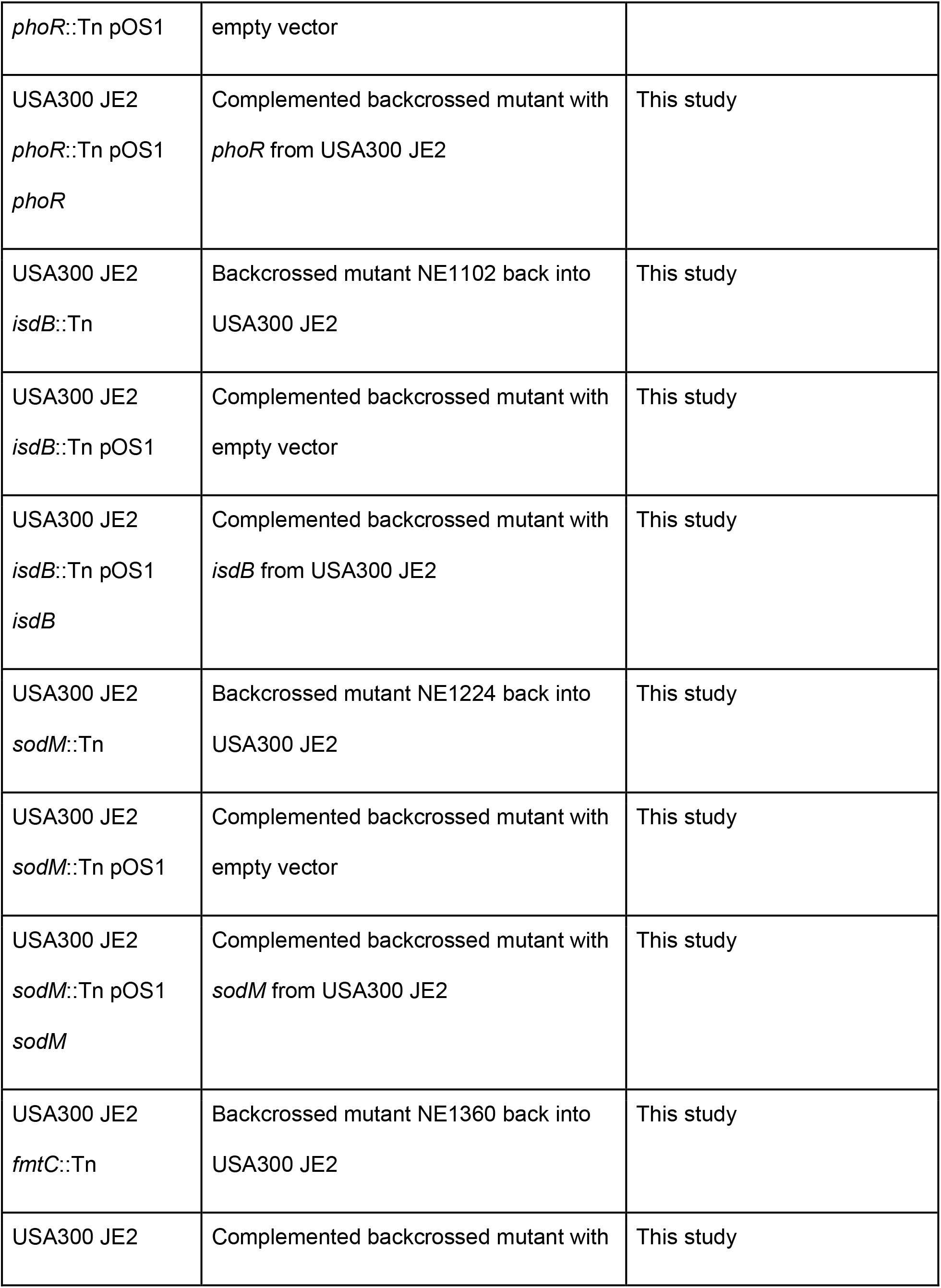
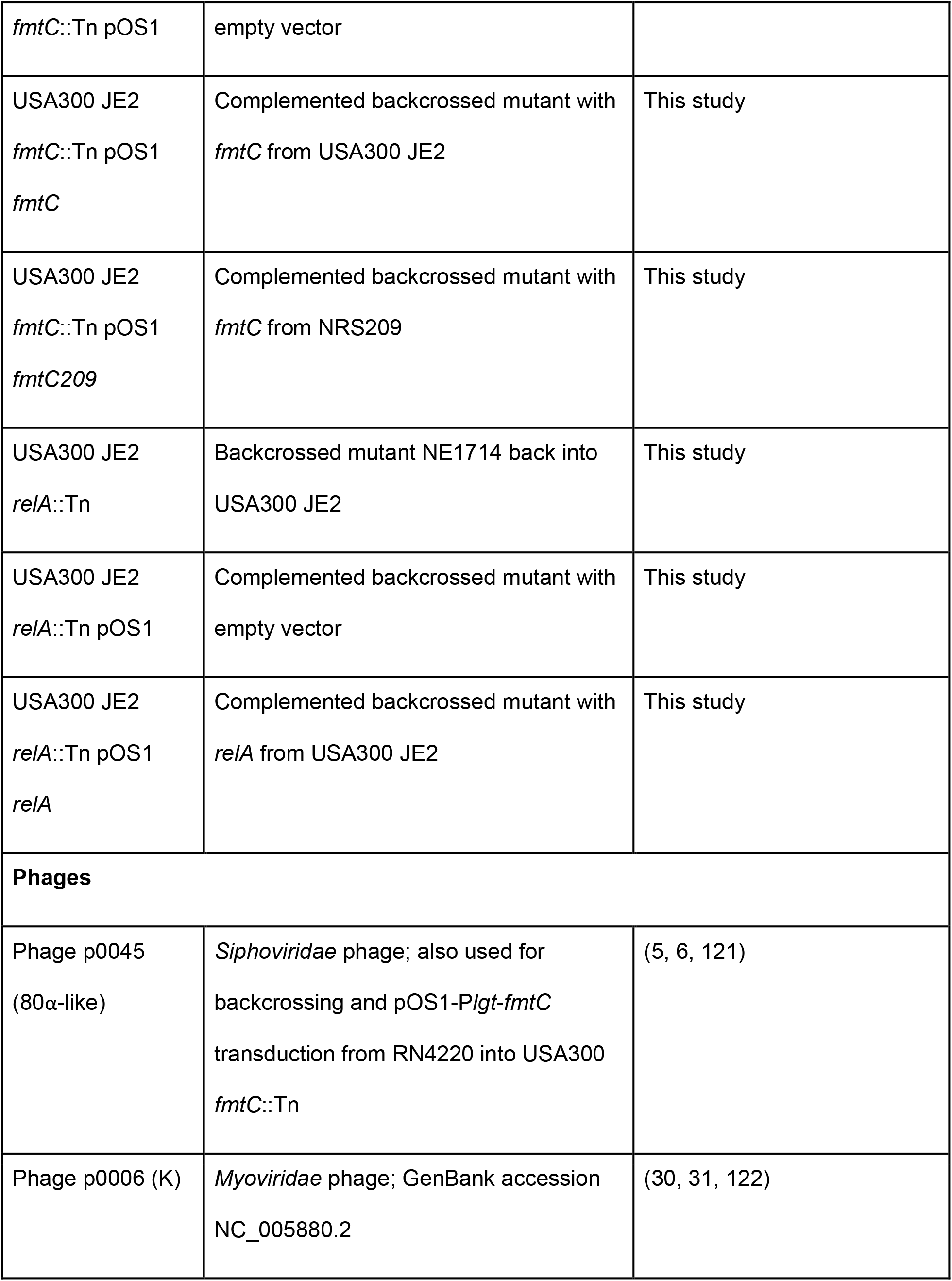
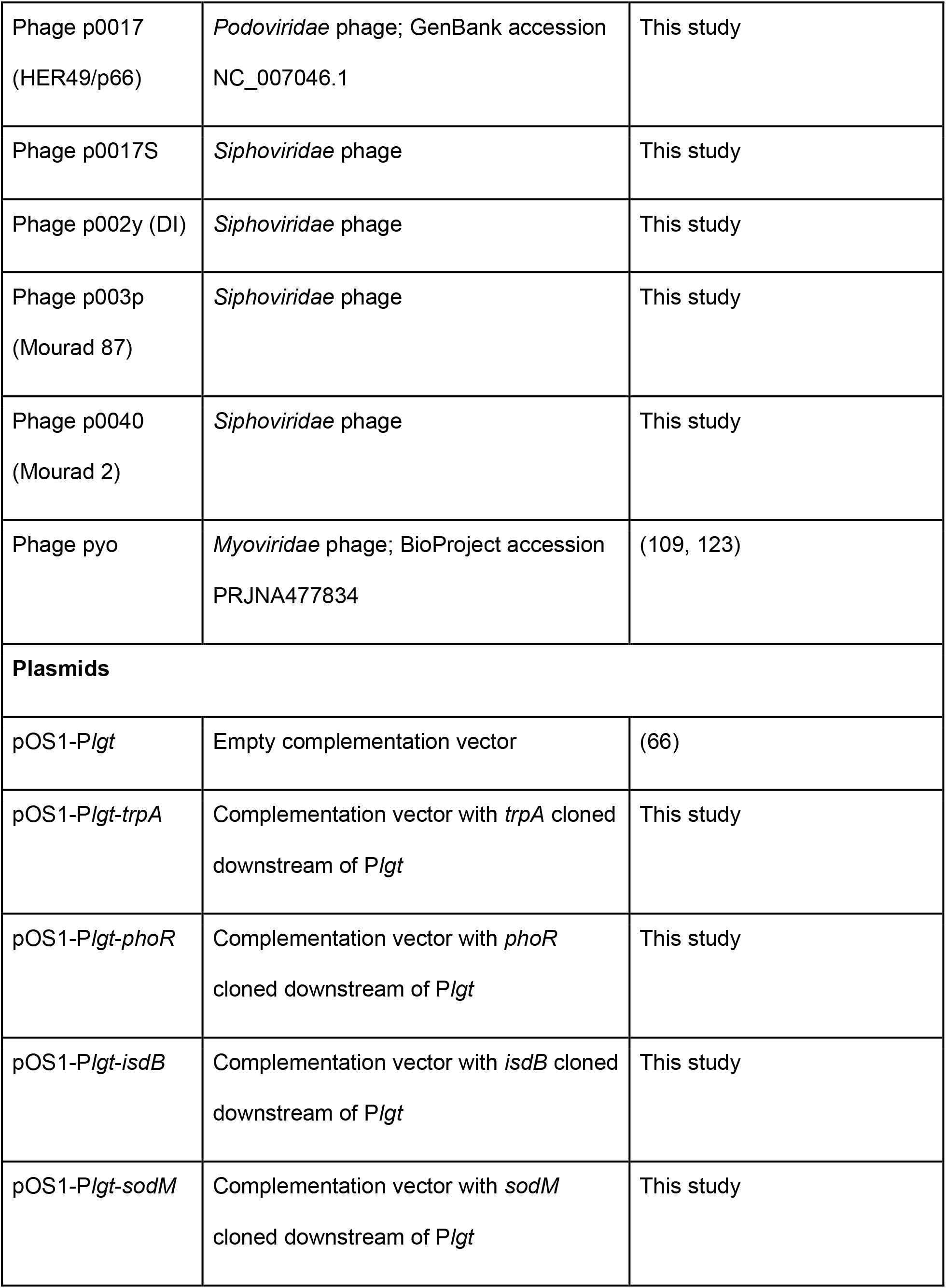
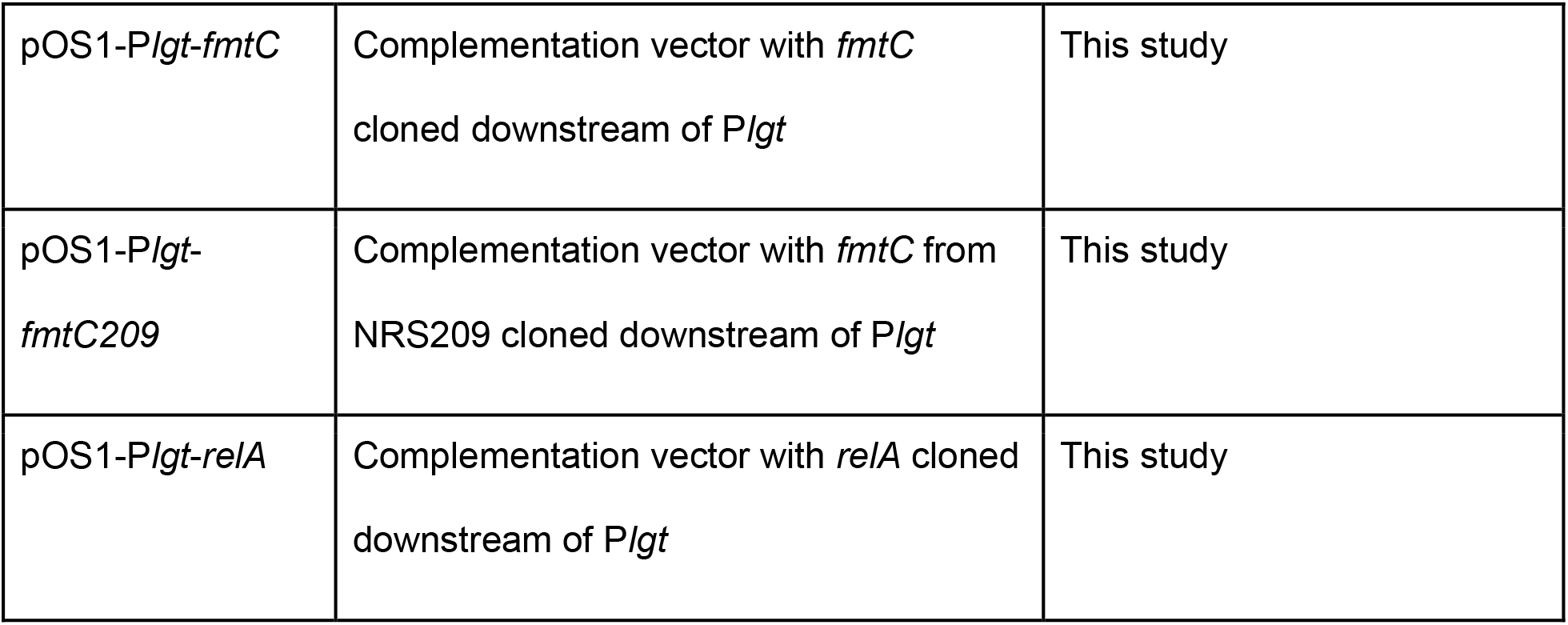
Strains, phages, and plasmids used for phage propagation and molecular genetic validation of GWAS results

Phages were propagated by inoculating a chunk of soft agar containing a plaque and surrounding bacteria into liquid medium. Phage lysates in TMG (Tris-magnesium-gelatin) buffer were spotted (4 μL) on a top agar (0.8% agar, 0.8% NaCl) lawn (5 mL) containing 0.2 mL of a 1:10 dilution of a RN4220 or RN4220 *tarM*::Tn overnight culture (18 hr growth, 37°C, 250 rpm). After overnight growth at 37°C, a chunk of soft agar containing a plaque and surrounding bacteria was inoculated in 35 mL of LB/TSB 2:1 with 5 mM CaCl_2_. This phage-bacterium co-culture was grown overnight at 37°C and 250 rpm, centrifuged for 20 minutes at 4,000 rpm, and filtered with a 0.45 μm syringe filter before being stored at 4°C. The resulting lysate was titered on RN4220 (*Siphoviridae* or *Myoviridae*) or RN4220 *tarM*::Tn (*Podoviridae*).

### Phage resistance/host range assays

259 previously genome-sequenced *S. aureus* strains consisting of 126 from the Network on Antimicrobial Resistance in Staphylococcus aureus (NARSA) repository (NCBI BioProject accession PRJNA289526) (39), 69 strains previously sequenced in a vancomycin-intermediate *S. aureus* (VISA) study (40) (PRJNA239001), and 64 strains previously sequenced in a cystic fibrosis (CF) lung colonization study (41) (PRJNA480016) were rapidly profiled for resistance to the eight phages using a high-throughput assay. Arrayed glycerol (50%) stocks of the strains were used to inoculate 96-well plates containing 200 μL of LB/TSB 2:1 with 5 mM CaCl_2_ in each well using a 96-pin replicator. Cultures were grown overnight at 37°C and 225 rpm. The following day, overnight cultures were diluted 1:10 in ddH_2_O. In order to permit phage adsorption, 10 μL of each phage lysate (~1e9 pfu/mL) was co-incubated with 10 μL of each overnight culture dilution for 30 minutes at room temperature in 96-well plates. 200 μL of molten LB/TSB/CaCl_2_ agar (LB/TSB 2:1 with 5 mM CaCl_2_ and 0.4% agar) was then added to each well containing the culture-phage mixtures and allowed to solidify. After incubation overnight (37°C), plates were photographed and final optical densities at 600 nm (OD_600_) per well measured using a plate reader (BioTek Eon). Strains were categorized as sensitive (0.1-0.4), semi-sensitive (0.4-0.7), or resistant (0.7 or greater) based on classifying average final OD_600_ from at least six replicates into three equal bins (with the third bin counting outlier resistant strains with OD_600_s above 1). Strains and host range phenotypes (quantitative and quantitative converted to ternary) are listed in Supplemental Tables S1 and S2.

High-throughput assays were also calibrated against a standard spot assay. 108 NARSA strains were tested for resistance to five of the eight phages listed previously (Phage p0045, p0006, p0017S, p002y, and p003p). Briefly, an overnight culture of each strain was diluted 1:10 in ddH_2_O and a top agar lawn (0.2 mL dilution per 5 mL molten top agar) was poured on a TSA plate. After solidification, each of the five lysates were spotted (4 μL) twice on the top agar lawn and let to dry. The plates were then incubated face up overnight at 37°C and the spots evaluated for clearing (sensitive), turbid clearing (semi-sensitive), or no clearing (resistant) the following day. High-throughput assay and spot assay phenotypes were compared in boxplots made with ggplot2 (42). The statistical significance of high-throughput assay phage resistance differences between all possible pairs of sensitive (S), semi-sensitive (SS), and resistant (R) strains were assessed with Wilcoxon signed-rank tests.

### Bioinformatic processing

Phage p0017 and pyo genomes were assembled from Oxford Nanopore reads with canu 2.0 (43). Hybrid Illumina/nanopore phage genome assemblies were constructed using Unicycler 0.4.8, filtering for contigs with coverage higher than 5x (44). Average nucleotide identity (ANI) was then determined amongst all phage contigs using fastANI 1.31 (45), which is shown as a lower-triangle identity matrix in **Supplemental Table S1**. All *S. aureus* genomes were processed using the Staphopia analysis pipeline (46), which included *de novo* assembly using SPAdes (47) and annotation using Prokka (48). The core-genome phylogenetic tree was constructed by first determining the core genome alignment for all tested strains with Roary (49), correcting for recombination with Gubbins (50), and then generating a maximum-likelihood phylogenetic tree with IQ-TREE (51). Strains (253 total) for which there are corresponding phage resistance phenotypes (quantitative and qualitative), BioProject, BioSample, and SRA accessions, sequence types, clonal complexes, isolation years, and isolation locations are listed in **Supplemental Table S2**. MLST (Multi-Locus Sequence Typing) Sequence types were identified for each genome with the mlst command line tool (52), which uses the PubMLST website (https://pubmlst.org/) (53). Quantitative phage resistance phenotypes were annotated on the tree using the Interactive Tree of Life (iTOL) (54).

### Preliminary phenotype analysis

Phage resistance phenotypes were initially placed on a core-genome phylogenetic tree and were associated with two factors - clonal complex (CC) and MRSA/MSSA genetic background. Phage resistance associations with CC and MRSA/MSSA were visualized in boxplots made with ggplot2 (42). Statistical significance of phage resistance differences between MRSA/MSSA was determined with Wilcoxon signed-rank tests. Statistical significance of overall phage resistance differences between represented CCs was determined using one-way analysis of variance (ANOVA) tests with or without phylogenetic correction.

### Measuring phylogenetic signal

Four different measures of phylogenetic signal were calculated for each phenotype - Abouheif’s C_mean_, Moran’s I, Pagel’s λ, and Blomberg’s K (55). Abouheif’s C_mean_ and Moran’s I were calculated with the abouheif.moran function from the adephylo R package (56) while Pagel’s λ and Blomberg’s K were calculated using the phylosig function from the phytools R package (57). Phylogenetic signal was determined using the core-genome phylogenetic tree annotated with quantitative phage resistance data previously described. Randomization tests for phylogenetic signal calculation were performed with 999 permutations of the data.

### Genome-wide association studies (GWAS)

Genotypes were associated with phage host range phenotype data using two different GWAS pipelines - pyseer 1.2.0 (58) and treeWAS 1.0 (59). Pyseer associated clusters of orthologous genes (COGs), core genome single nucleotide polymorphisms (SNPs), and k-mers between lengths 6 and 610 with each phenotype, while treeWAS only associated biallelic core genome SNPs with the phenotype. TreeWAS used the recombination-corrected core-genome phylogeny for population structure correction while pyseer used a conversion of the phylogeny into a kinship matrix. The core genome alignment was rearranged to set N315 as the reference (first sequence). We chose N315 as reference because it was used as a global *S. aureus* reference for the Staphopia project (46). SNPs were called from the core genome alignment with snp-sites (60). For identifying significantly associated genetic determinants, a Bonferroni correction of 0.05/6,058 or 8.25e-6 was set for COG GWAS, 0.05/15,557 or 3.21398e-6 for SNP GWAS, and 0.05/2,304,257 or 2.17e-8 for k-mer GWAS, counting the numbers of intermediate-frequency COGs, biallelic core genome SNPs, and unique k-mers as hypotheses to be tested, respectively.

Pyseer SNP and COG association analyses performed multidimensional scaling (MDS) on a mash distance matrix between tested strains to correct for population structure. Pyseer SNP association was performed with a fixed effect (for variant and covariate lineage) model, the default 10 multidimensional scaling (MDS) dimensions retained, and lineage effect testing on each quantitative phage resistance/host range phenotype for all biallelic core genome SNPs. Pyseer COG association was performed with a fixed effects model on each phenotype and 9 MDS dimensions retained for intermediate frequency COGs (**Supplemental Figure S1**). Pyseer k-mer association was performed with a FaST-LMM linear mixed (combined fixed variant/covariate lineage and random kinship effects) model on each quantitative phenotype for unique k-mers between 6 and 610 bp in length extracted from genomes of all tested strains. Pyseer k-mer association analyses used a kinship matrix between tested strains constructed from the core-genome phylogeny to correct for population structure and set a minor allele frequency cutoff for analysis of 1%, like SNP and COG analyses. SNP and k-mer association p-values were visualized relative to genetic coordinates using Manhattan plots (with phandango) (61). Associations for all k-mers were assessed for p-value inflation (exceeding the observed/expected p-value diagonal below 1e-2) using Q-Q plots (**Supplemental Figure S2**). Significant SNPs and k-mers were annotated using SnpEff (62) (relative to the Roary N315 core genome sequence) and downstream analysis scripts included with pyseer, respectively, identifying the genes containing the genetic elements (or near the genetic elements, in the case of k-mers) and mutation effects, in the case of SNPs.

TreeWAS was performed for each phage resistance phenotype using the R package with core genome alignment, IQ-TREE core-genome phylogeny, and quantitative phage resistance phenotype as inputs and with default parameters. Significant treeWAS SNPs were annotated using SnpEff (62) relative to the core genome sequence of strain N315 (63).

### Functional annotation and network analysis of significantly associated genes

Genes with significant association from the GWAS study (containing SNPs, and either near or overlapping with k-mers) were then used to identify enriched protein functions or possible protein-protein interactions. Gene name lists for each phage were converted to NCTC 8325 RefSeq protein accession lists for use with STRING (64) and PANTHER (65), which depend on NCTC 8325 *S. aureus* accessions. To convert genes containing significant SNPs to NCTC 8325 accessions, Roary N315 core genes were aligned against NCTC 8325 RefSeq proteins with NCBI blastx (1 maximum target sequence, 1 maximum high scoring pair, default e-value). Gene names matching NCTC 8325 RefSeq accessions were converted for each significant SNP using these alignment results. To convert genes containing significant k-mers to NCTC 8325 accessions, all significant genes were aligned against NCTC 8325 RefSeq proteins with blastx (1 maximum target sequence, 1 maximum high scoring pair, default e-value). Gene names matching NCTC 8325 RefSeq accessions were converted for each significant k-mer using these alignment results. Any gene names not mapped to any NCTC 8325 RefSeq protein accessions after this procedure were left unchanged. Lists of significant genes for each phage, for all phage morphological classes (*Siphoviridae*, *Myoviridae*, and *Podoviridae*), and for each life cycle type (virulent or temperate) were used as inputs for STRING and PANTHER. STRING network properties (nodes, edges, average node degree, average local clustering coefficient, expected number of edges, and PPI enrichment p-value) were saved for each input, while PANTHER functional classification and statistical overrepresentation test analyses were performed for each input with respect to molecular function, biological process, cellular component, protein class, and pathway.

### Genetic validation of novel phage resistance mechanisms

Six genes (*trpA, phoR, isdB, sodM, fmtC*, and *relA*) found to contain significantly associated SNPs or k-mers for any phage resistance phenotype were validated to cause phage resistance changes when knocked out in a single *S. aureus* genetic background (USA300 JE2). Transposon insertion mutants in each gene were selected from the Nebraska Transposon Mutant Library (NTML) (37) and backcrossed into USA300 JE2 through the transduction method previously described (38) to eliminate any possible secondary acquired mutations. Backcrossed mutants were then complemented with each gene cloned into the vector pOS1-P*lgt* (66). Relevant strains (selected mutants and complemented strains) are listed in **Table 1**. Growth curves were performed on all listed strains (**Supplemental Figure S4**). USA300 JE2, respective transposon mutants, empty vector controls, or complemented mutants were inoculated with a 96-pin replicator from arrayed frozen glycerol stocks into 96-well plates containing 200 μL LB/TSB 2:1 with 5 mM CaCl_2_ or the same medium supplemented with 10 μg/mL chloramphenicol in each well. We then diluted each culture 1:100 in fresh LB/TSB 2:1 with 5 mM CaCl_2_ or the same medium supplemented with 10 μg/mL chloramphenicol and collected growth curves on a BioTek Eon plate reader (37°C, 225 rpm agitation, OD_600_ measured every 10 minutes).

Genes were cloned into pOS1-P*lgt* either through splicing overlap extension (SOE) PCR (*trpA, phoR*, and *sodM*) or through NEB HiFi assembly (*isdB, fmtC*, and *relA*). Each gene and pOS1-P*lgt* were amplified with the primers listed in **Supplemental Table S3** to create overlap into the corresponding fragment using NEB Q5 High-Fidelity DNA Polymerase according to manufacturer’s directions. All genes were amplified from USA300 JE2 genomic DNA except for fmtC, which was amplified both from USA300 JE2 and NRS209. Genes were cloned into the same site downstream of the P*lgt* promoter. For SOE PCR, AMpure XP bead-purified gene and vector fragments were mixed together at a ratio of 1:59 and amplified for 20 cycles with NEB Q5 High-Fidelity polymerase at an annealing temperature of 60°C. For HiFi assembly, purified gene and vector fragments were mixed together at a ratio of 1:2 (less than 0.2 pmol DNA total) and incubated with NEBuilder HiFi DNA Assembly Master Mix for 3 hours at 50°C. SOE PCR and HiFi assembly products were transformed into NEB DH5α competent cells (High Efficiency), plated on LB agar with ampicillin (100 μg/mL), and grown overnight at 37°C. Transformants were verified by colony PCR with respective LF and RR primers listed in **Supplemental Table S3**. Plasmids were extracted from verified transformant overnight cultures with the Promega PureYield Plasmid Miniprep System. These plasmids were then transformed into *E. coli* IM08B (67) to improve electroporation efficiency into the USA300 JE2 transposon mutants.

Electrocompetent *S. aureus* cells (USA300 JE2 transposon mutants) were prepared as previously described (68). *S. aureus* electrocompetent cells were electroporated with 2 μg of ethanol-precipitated plasmid DNA (empty vector and vector with insert corresponding to transposon insertion). Electrocompetent cells were first thawed, centrifuged, and resuspended in 50 μL 10% glycerol/0.5 M sucrose. After adding plasmid DNA, cells were transferred to 0.1 cm electroporation cuvettes and pulsed at 2.1 kV, 100 Ω, and 25 μF. Immediately after electroporation, 1 mL of TSB/0.5 M sucrose was added to the cuvette and the culture was transferred to an Eppendorf tube to recover for 90 minutes at 37°C and 250 rpm. Dilutions of the outgrowth were plated on TSA with chloramphenicol (10 μg/mL) and grown overnight at 37°C. Electroporants were verified by colony PCR with respective LF and RR primers listed in **Supplemental Table S3**.

pOS1 *fmtC* and *relA* were introduced into USA300 JE2 transposon mutants, however, by transduction from RN4220. *S. aureus* RN4220 was electroporated with pOS1 *fmtC* (USA300), pOS1 *fmtC* (NRS209), and pOS1 *relA* plasmids according to the procedure described previously. Plasmids were then transduced from RN4220 to USA300 JE2 transposon mutants according to a procedure previously published (38). Briefly, a recipient strain was infected with donor phage at a MOI of 0.1 after supplementing with CaCl_2_. The infected culture was then outgrown in TSB supplemented with sodium citrate to prevent phage lysogeny. The outgrowth culture was plated on TSA supplemented with both chloramphenicol (10 μg/mL) and sodium citrate (40 mM) to select for plasmids and inhibit lysogeny, respectively.

Mutants and their complemented derivatives were assessed for phage resistance and host range both through the high-throughput assay described previously and the efficiency of plating (EOP) assay (69) to assess bacterial growth in the presence of phage and phage plaquing efficiency, respectively. The high-throughput host range assay was performed as described earlier, but strain overnight cultures were grown in LB/TSB 2:1 with 5 mM CaCl_2_ supplemented with chloramphenicol (10 μg/mL) to maintain plasmid selection in the case of complemented strains for this and the EOP assay. The EOP assay was performed by spotting 4 μL of neat through 1e-8 dilutions of phages p0045, p0006, p0017, p0017S, p002y, p003p, p0040, and pyo on lawns (0.2 mL of a 1:10 overnight culture dilution mixed with 5 mL of top agar) of a test and reference (USA300) strain. Lawns were poured on TSA plates. EOP was calculated by dividing phage titer on the test strain by that on the reference strain.

Additional experiments on the *trpA* mutant set and phage p003p examined bacterial survival after performing the phage/culture soft agar coincubation of the high-throughput assay. The high-throughput assay was performed as described earlier for six replicates of USA300, USA300 *trpA*::Tn, USA300 *trpA*::Tn pOS1, and USA300 *trpA*::Tn pOS1 *trpA* strains. Corresponding ODs were recorded as described for the high-throughput phage host range assay (**Supplemental Figure S5A**). Agar plugs were then removed with toothpicks, placed in 0.8 mL volumes of sterile TMG, and broken apart by vortexing. The resuspensions were then serially diluted in TMG and 4 uL of 1e-1 through 1e-6 dilutions were spotted four times on TSA plates. Dilution plates were grown overnight at 37°C and colonies counted the following day to determine surviving CFU in each condition (**Supplemental Figure S5B**).

### Construction of phage resistance phenotype predictive models

Phage resistance predictive models were constructed using three methods - random (decision) forests, gradient-boosted decision trees, and neural networks. Random forests were generated using the randomForest R package, gradient-boosted decision trees were generated with the XGBoost R package (70). Ternary (S, SS, or R) phenotypes converted from the original high-throughput assay quantitative phenotypes (described in the phage host range assay methods) were set as the response variable, while either presence/absence of each significant genetic element, each k-mer, or one of the previous two sets (all elements or just k-mers) and both strain sequence type (ST) and clonal complex (CC) were set as predictor variables. Random forest and XGBoost predictive accuracy and receiver operating characteristic (ROC) area under the curve (AUC) was determined on the validation set through multiple replicates of 10-fold cross-validation, in which alternating tenths of the data are used for validation while the model is trained on the remaining data. The optimal number of rounds (iterations) for XGBoost was determined for each phage and set of input predictor variables with 5-fold cross-validation. XGBoost model training also used the softmax objective for multiclass (three classes - S, SS, and R) classification.

Neural network model construction was more complicated as it involved a preprocessing step to balance datasets where necessary. Oversampling or a combination of over- and undersampling methods was performed to balance specific datasets. For the oversampling method, new samples of the minority classes were randomly generated with replacement so that the number of samples for each class would be equal to that of the majority class in the original dataset. For the combination method, the Synthetic Minority Over-sampling Technique (SMOTE) for over-sampling and Tomek links for under-sampling were performed together. However, for phages with limited cases for one class type, such as p002y, we cannot conduct undersampling. Therefore, for such datasets, only the oversampling method was performed. Then the new balanced datasets were split into training and validation sets with 30% validation. Random splits were performed four times to generate four replicates for evaluation, each with different train and test datasets. Each replicate was evaluated as before with validation set prediction accuracy and ROC AUC.

Neural network models were constructed three ways - 1) with or without oversampling or an over-/undersampling combination alone, 2) also with a regularizer and dropout layer, or 3) also with lasso regression for feature selection. All methods use ADAM (71) for optimizing and sparse categorical cross-entropy for loss. For imbalanced datasets, the oversampling and combination over/undersampling methods were used as well, if possible. The fully connected neural network was constructed based on the selected, balanced dataset. We then found both training and prediction accuracy to evaluate performance for each network model. We note that network models were optimized for each replicate training set, which means there may be different network models for the four replicates. In the first method, fully connected neural network models were constructed on datasets either originally balanced or balanced after over-/combination methods, with no further correction. Since some network models have high prediction accuracies, it is possible that these models are overfitting, so the second method adds a regularizer and a dropout layer to fully connected neural networks as new models. Finally, for some network models, the prediction accuracies were not as high as others. Thus, in the third method, lasso regression was performed to select important features and improve performance. A neural network model was constructed on the new dataset based on these selected features.

Information entropy was compared to average randomForest and XGBoost 10-fold cross-validation and neural network predictive accuracies and ROC AUCs after calculation through using the following equation (72), where P_x_(x_i_) is the probability of event xi, and the three possible events are S, SS, and R phenotypes:

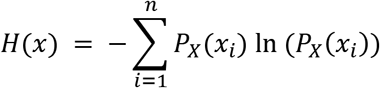

## Results

### Development of a novel high-throughput host range assay

In order to evaluate host range on a large number of *S. aureus* strains in a quantitative manner, we developed a high-throughput host range assay (**Figure 1**) described in the Materials and Methods section. This assay measures the extent that phages cause retardation of growth compared to a control. Before using data from the high-throughput assay for further analysis, we calibrated it against the traditional spot assay (**Figure 1A**), which measures whether phages cause lysis in a lawn of bacterial cells. We compared spot assay results (sensitive - S, semi-sensitive - SS, or resistant - R) for 108 strains and five phages to the strains’ average final soft agar turbidity (OD_600_) in the high-throughput assay (**Figure 1B**). For all phages tested, turbidity was significantly higher (p<0.05, Wilcoxon signed-rank test) for spot-resistant strains relative to spot-sensitive strains. For all phages tested but p003p, the turbidity was significantly higher for spot-semi-sensitive strains relative to spot-sensitive strains. However, for only phages p0006 and p003p were turbidities significantly higher for spot-resistant strains relative to spot-semi-sensitive strains. Thus, for all phages but p003p, it was possible to tell spot-sensitive from spot-semisensitive strains by the high-throughput assay, but only for phages p0006 and p003p was it possible to tell spot-semi-sensitive from spot-resistant by the new assay. Overall, these results showed strong agreement between the lysis-based spot assay and the high-throughput growth-based assay for differentiating between sensitive and resistant/semi-sensitive phenotypes.

**Figure 1:**
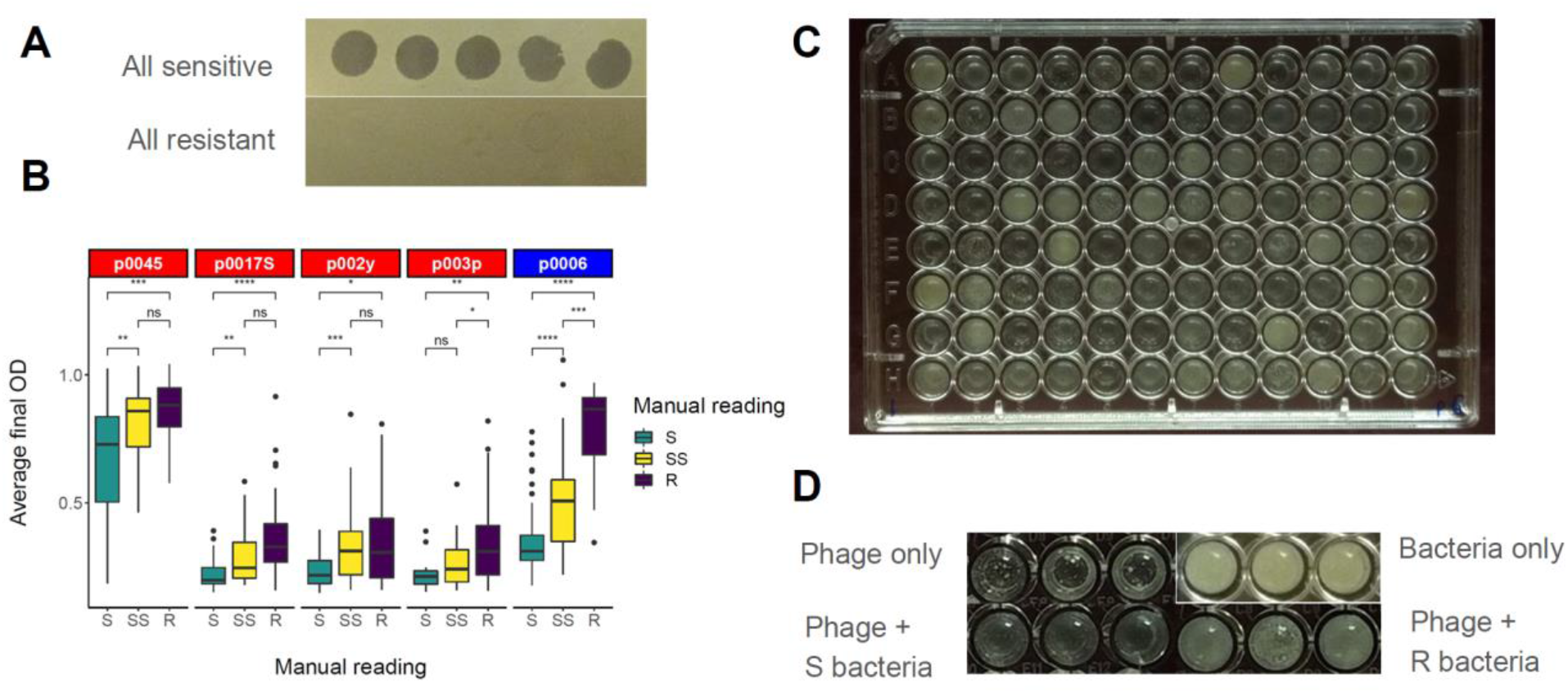
Development of the high-throughput phage host range assay. A) Example of fully sensitive (NRS149) and fully resistant (NRS148) spot assay phenotypes for five test phages (p0045, p0006, p0017S, p002y, and p003p). *Siphoviridae* are listed in red, *Myoviridae* in blue, and *Podoviridae* in purple. B) Calibration of the high-throughput assay against qualitative spot assay phenotypes (S - sensitive, complete clearing; SS - semi-sensitive, cloudy clearing; R - resistant, no clearing) collected with the spot assay for 108 NARSA strains and the five phages listed in A. Data represent the distribution of average high-throughput assay measurements for strains evaluated as S, SS, or R in corresponding spot assays. Wilcoxon signed-rank test significance values for each possible comparison are listed at the top of the corresponding boxplots (ns - not significant, * - 0.01 to 0.05, ** - 0.001 to 0.01, *** - 0.0001 to 0.001, **** - less than 0.0001). C) Example high-throughput assay results from one 96-well plate containing 96 NARSA strain overnight cultures co-incubated with phage p0006 (K). D) Example high-throughput assay phenotypes for a sensitive *S. aureus* strain, resistant strain, bacteria without phage, and phage without bacteria.

### Host range is associated with clonal complex but not methicillin resistance

We evaluated the host range of eight phages belonging to the *Siphoviridae*, *Myoviridae*, and *Podoviridae*. *Siphoviridae* (p0045, p0017S, p002y, p003p, and p0040), *Myoviridae* (p0006 and pyo), and *Podoviridae* (p0017) were most closely related to others of the same class, but not related at all to those of other classes (**Supplemental Table S1**). Amongst the *Siphoviridae*, p003p was the most divergent from the others (between 97.75 and 97.83% similar to the others). On the host side, host range was determined on a set of 259 *S. aureus* strains representing 47 already-defined sequence types (STs) and 17 already-defined clonal complexes (CCs) against eight phages (253 strains with sequence data are included in **Supplemental Table S2**). The most common STs were 5 (25.69%), 8 (13.04%), 30 (6.72%), 105 (4.35%), and 121 (3.16%), while the most common CCs were 5 (37.15%), 8 (23.32%), 30 (12.25%), 121 (5.14%), and 1 (4.74%), respectively. The most common strain isolation years were 2005 (31.92%), 2012 (14.08%), 2002 (12.68%), 2017 (7.51%), and 2018 (7.04%), while the most common isolation locations were the United States (61.26%), France (19.76%), the United Kingdom (11.46%), and Japan (1.19%). Strain isolation years ranged from 1935 to 2018.

Phages p0045, p0040, the two temperate phages, and p0017, the sole tested podovirus, had the highest proportions of resistant strains (71.8, 38.2, and 35.9%, respectively) amongst those tested (**Figure 2A** and **Table 2**). The average and median final turbidities amongst tested strains were likewise highest for these phages (0.80/0.88, 0.61/0.60, and 0.56/0.54, for p0045, p0040, and p0017, respectively). On the other hand, phages p0017S, p002y, p003p, and pyo, all virulent *Sipho*- or *Myoviridae*, had the lowest proportions of resistant strains (0.8, 1.2, 1.2, and 1.5%, respectively) and average/median final turbidities (0.31/0.27, 0.27/0.22, 0.32/0.31, and 0.26/0.21, respectively). Phage p0006 had an intermediate proportion of resistant strains (15.4%) and average/median final turbidity (0.49/0.44). Strains were resistant to between zero and six phages (**Figure 2B**), with a median of two. The strains NRS148, NRS209, and NRS255 were resistant to six phages, the most among any strains. Phage host ranges were most similar (concordant - defined by number of strains with identical phenotypes between two phages) between phages p0017S, p002y, p003p, and pyo, but least similar between phage p0045 and the previous set of four phages (**Figure 2C**).

**Figure 2:**
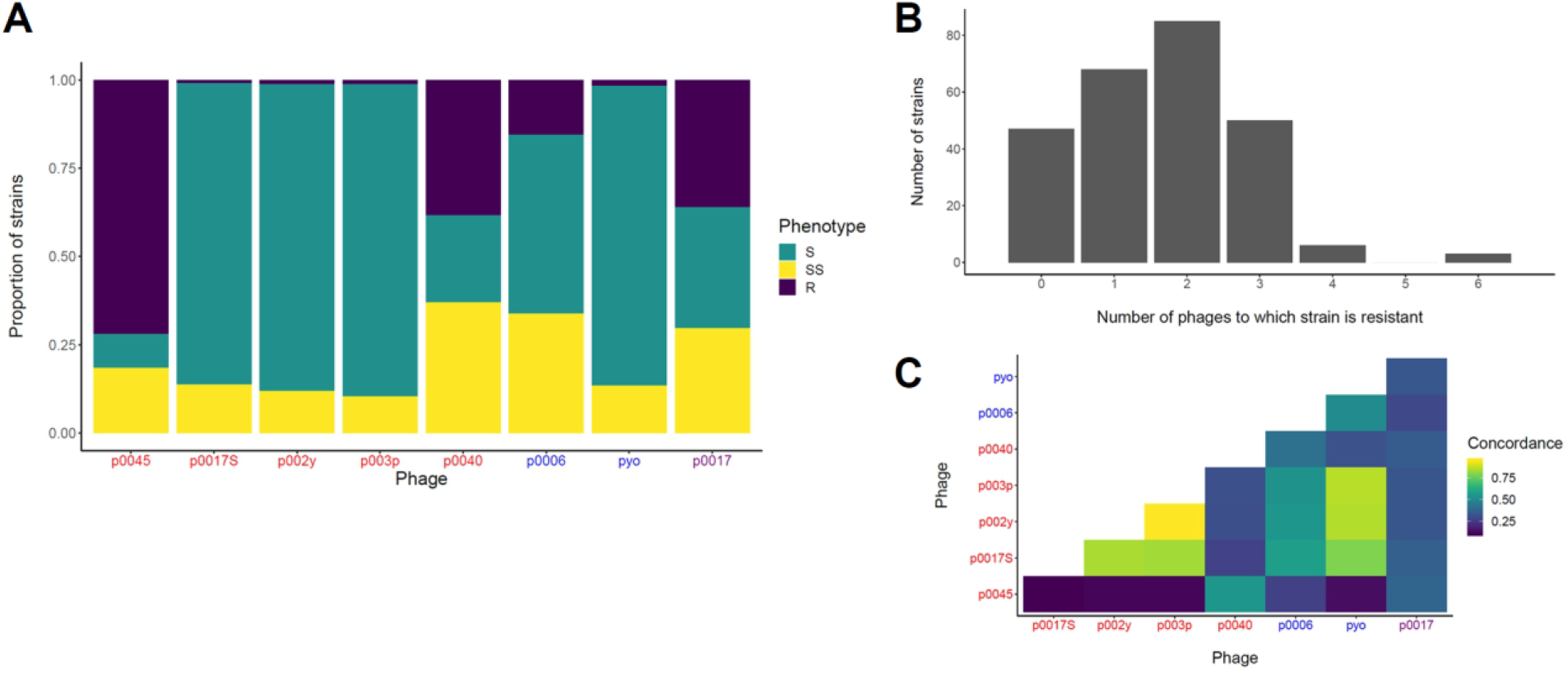
Host range distribution, concordance, and multiple phage resistance. A) Number of strains that fall into host range categories for each phage. Sensitive (S) corresponds to an OD_600_ of 0.1-0.4, semi-sensitive corresponds to an OD_600_ of 0.4-0.7, and resistant corresponds to an OD_600_ of 0.7 or higher. B) Histogram of number of phages to which strains are resistant, by the previous definition. C) Concordance matrix of the tested phages’ host ranges. Concordance is defined as the number of strains with identical phenotypes between two phages. *Siphoviridae* are listed in red, *Myoviridae* in blue, and *Podoviridae* in purple.

**Table 2:** Summary statistics of phage host range phenotypes. For each phage, the number of strains falling into each category were counted. These phenotypes were determined for each phage using the high-throughput assay. Sensitive (0.1-0.4), semisensitive (0.4-0.7), and resistant (0.7 and higher) strain numbers and percentages are listed first, followed by mean, standard deviation, and median quantitative phenotypes for all tested strains. Statistics summarize at least six biological replicates for each page.

| Phage | p0045 | p0006 | p0017 | p0017S | p002y | p003p | p0040 | pyo |
| --- | --- | --- | --- | --- | --- | --- | --- | --- |
| Sensitive (%) | 25 (9.7%) | 131 (50.6%) | 89 (34.4%) | 221 (85.3%) | 225 (86.9%) | 229 (88.4%) | 64 (24.7%) | 220 (84.9%) |
| Semi-sensitive (%) | 48 (18.5%) | 88 (34.0%) | 77 (29.7%) | 36 (13.9%) | 31 (12.0%) | 27 (10.4%) | 96 (37.1%) | 35 (13.5%) |
| Resistant (%) | 186 (71.8%) | 40 (15.4%) | 93 (35.9%) | 2 (0.7%) | 3 (1.2%) | 3 (1.2%) | 99 (38.2%) | 4 (1.5%) |
| Mean | 0.80 | 0.49 | 0.56 | 0.31 | 0.27 | 0.32 | 0.61 | 0.26 |
| Stdev | 0.24 | 0.20 | 0.27 | 0.12 | 0.12 | 0.12 | 0.23 | 0.14 |
| Median | 0.88 | 0.44 | 0.54 | 0.27 | 0.22 | 0.31 | 0.60 | 0.21 |

We also examined whether there were significant associations between clonal complex (CC) or methicillin resistant *S. aureus* (MRSA) genetic background and each phage host range phenotype (**Figure 3**). We hypothesized CC would correlate with host range given that type I restriction-modification specificity is strongly associated with CC (73, 74), restricting the infection of a strain by phage propagated in a strain of a different CC. We hypothesized MRSA genetic background may also affect host range, given that the phage receptor WTA is required for methicillin resistance (75) but MRSA strains can tolerate more defects in WTA biosynthesis than MSSA strains (76). However, MRSA/MSSA phenotypic differences were only significant for phage 0040 (p<0.001, Wilcoxon signed-rank test; **Figure 3B**). There were significant differences in phage resistance between individual CCs for all phages (p<0.05, Tukey Honest Significant Differences based on one-way ANOVA; **Supplemental Table S4**) and significant overall differences amongst all CCs (one-way ANOVA) for all phages (p<0.05). Overall, these results indicate MRSA genetic background for the most part is not associated with the host range of these phages, while CC overall affects all tested phages’ host ranges.

**Figure 3:**
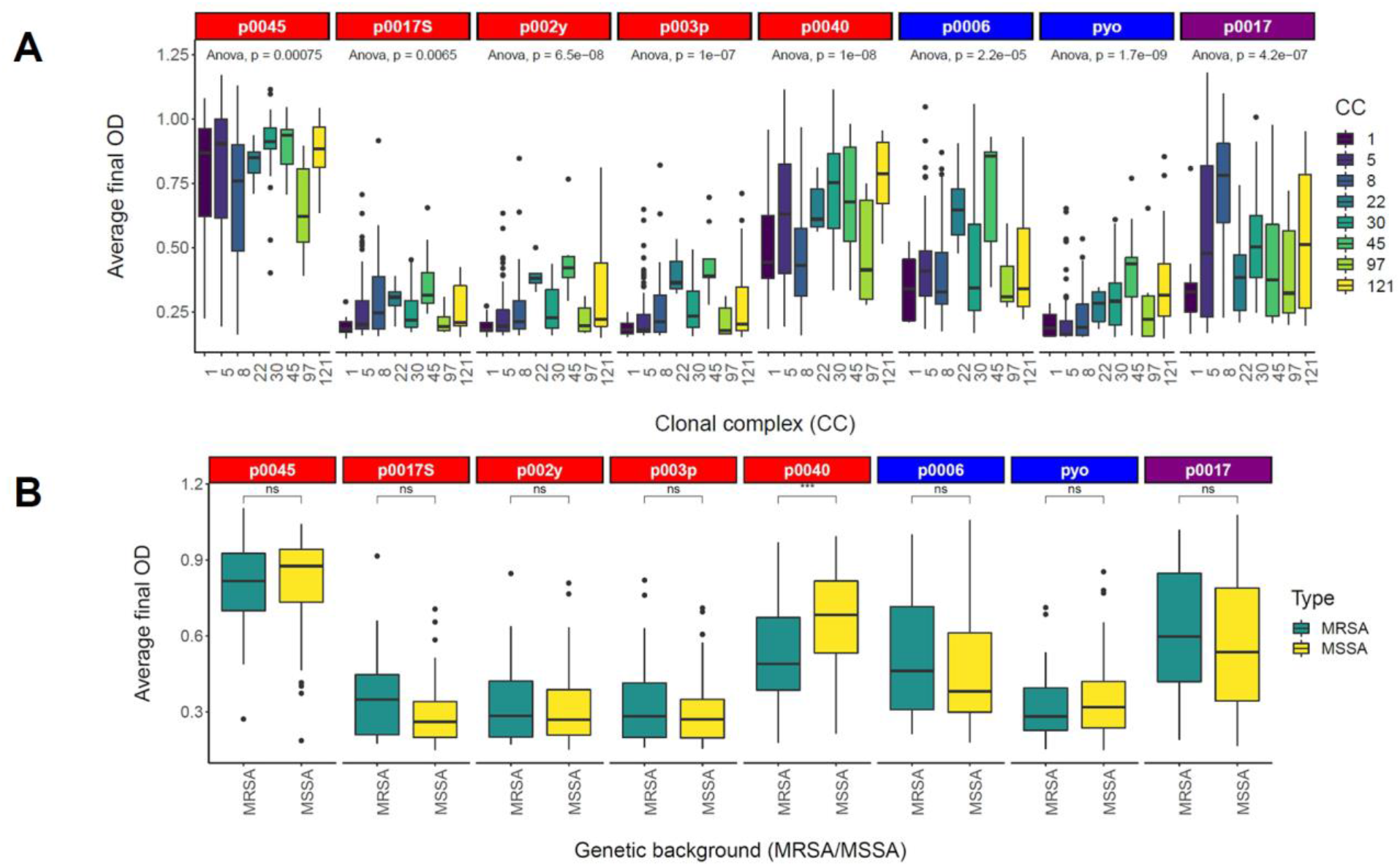
Phage resistance is related to clonal complex (CC) but not MRSA genetic background. Data represent the distribution of average high-throughput assay measurements for strains belonging to each presented CC (all 259 strains) (A) or MRSA/MSSA (126 NARSA strains) (B) genetic background. One-way ANOVA significance values for overall differences amongst CCs presented and Wilcoxon signed-rank test significance values for MRSA/MSSA differences are listed at the top of the corresponding boxplots (ns - not significant, * - 0.01 to 0.05, ** - 0.001 to 0.01, *** - 0.0001 to 0.001, **** - less than 0.0001). *Siphoviridae* are listed in red, *Myoviridae* in blue, and *Podoviridae* in purple.

Resistance to each phage is highly homoplasious, emerging independently in multiple CCs (**Figure 4**). We estimated phylogenetic signal by calculating Moran’s I, Abouheif’s C_mean_, Pagel’s λ, and Blomberg’s K (55) for each phage host range phenotype, which resulted in statistically significant values in every case (**Table 3**). Both Moran’s I and Abouheif’s C_mean_ values fell between 0.17 and 0.37. Pagel’s λ values all were nearly 1, while Blomberg’s K values approached 0. Pagel’s λ values around 1 and Moran’s I/Abouheif’s C_mean_ values around 0 support a Brownian motion model (the phylogeny structure alone best explains the trait distribution), but Blomberg’s K values around 0 suggest trait variance at the tips is greater than that predicted by the phylogeny under a Brownian motion model. All calculated phylogenetic signal values were statistically significant (p<0.05 for randomization tests based on 999 simulations). Taken together, these results suggest the structure of the phylogeny could explain the host ranges of the tested phage as expected under a Brownian motion model (random distribution of phenotypes amongst strains directed by the phylogeny overall). This neutral phylogenetic signal agrees with the previous finding that CC is associated with host range (**Figure 3A** and **Supplemental Table S4**). While there is a CC association with host range, strain-specific effects may be even stronger than CC-specific effects, resulting in weak net phylogenetic signals.

**Figure 4:**
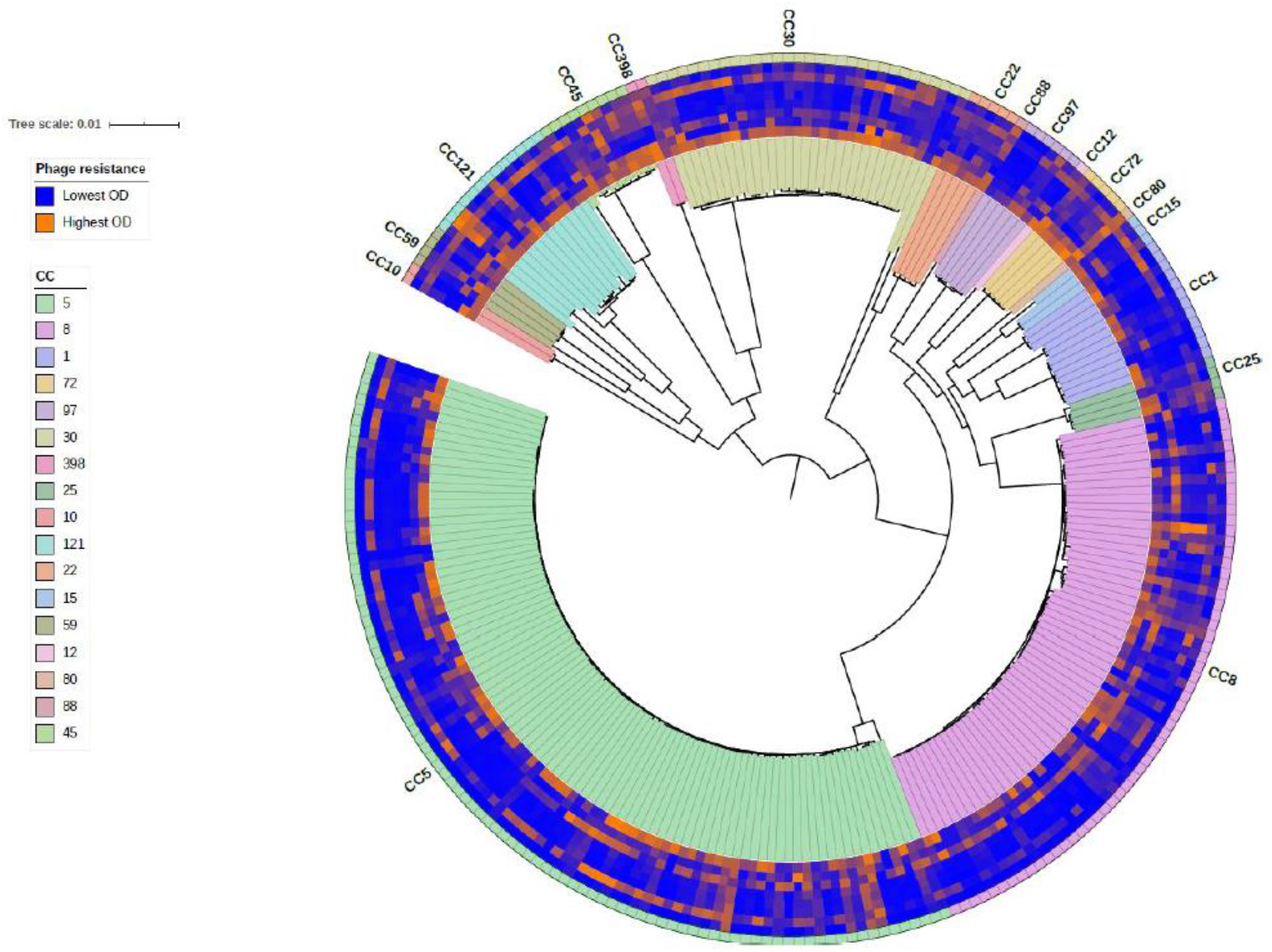
Phage resistance across the *S. aureus* species. Average high-throughput phage host range assay phenotypes (of at least six replicates) and corresponding strain clonal complexes were placed on a maximum-likelihood, midpoint-rooted core-genome phylogeny and visualized with the Interactive Tree of Life (iTOL) (54). Phenotypes are presented on a scale from blue (lowest OD_600_, most sensitive) to orange (highest OD_600_, most resistant). Phenotypes from inside to outside correspond to phages p0045, p0006, p0017, p0017S, p002y, p003p, p0040, and pyo. CCs are shaded inside and outside the circumference of the tree.

**Table 3:** Measures of phylogenetic signal for each phage resistance phenotype. Values that are significant are shown in bold. Significance was determined for 999 random permutations of the data.

| Phage | Moran's I | Abouheif's $C_{\text{mean}}$ | Pagel's $\lambda$ | Blomberg's K |
| --- | --- | --- | --- | --- |
| p0045 | **0.23** | **0.23** | **1.00** | **0.005** |
| p0006 | **0.17** | **0.17** | **1.00** | **0.008** |
| p0017S | **0.32** | **0.32** | **1.00** | **0.007** |
| p002y | **0.23** | **0.23** | **1.00** | **0.008** |
| p003p | **0.30** | **0.30** | **1.00** | **0.012** |
| p0040 | **0.28** | **0.28** | **1.00** | **0.014** |
| pyo | **0.36** | **0.37** | **1.00** | **0.006** |
| p0017 | **0.31** | **0.31** | **1.00** | **0.014** |

### GWAS reveals novel genetic determinants of host range

We used the GWAS tools pyseer (58) and treeWAS (59) to identify genetic loci strongly associated with the phage host range phenotype (**Supplemental Figure 3A, Table 4**). We chose these tools because they represent two alternatives for population structure correction - identifying principal components of a distance matrix (pyseer) and testing against phenotypes simulated based on the phylogeny (treeWAS). pyseer identified COGs, SNPs, and k-mers beyond the respective multiple-corrected significance thresholds in all phages. Most phages lacked k-mer p-value inflation with the exceptions of p0017S, p002y, and p003p, based on associated Q-Q plots (scatter above the diagonal at p-values of 1e-2 or more indicated p-value inflation; **Supplemental Figure S2**). The number of significant COGs detected ranged from 48 (p0017S) to 347 (pyo). Significant SNPs were detected for all phages but p0045 and p0017S and ranged from 1 (p0017) to 249 (pyo). Significant SNPs were identified in *tarJ* (pyo - 672A>G synonymous) and *tagH* (p002y - 848T>C missense and 873A>T missense; pyo - 848T>C missense, 873A>T missense, and 876C>T synonymous). TarJ is responsible for activating ribitol phosphate with CTP to form CDP-ribitol (77), while TagH is a component of the ABC transporter that exports WTA to the cell surface (9). A substantial number of the significant p0017 k-mers (1,382, −log(p-value) = 12.259) mapped to the recently discovered host range factor *tarP*. TarP was shown to confer podovirus resistance by transferring N-acetylglucosamine to the C3 position of ribitol phosphate (14). Significant k-mers also mapped to *hsdS* (32 for p002y, −log(p-value) = 9.33; 6 for p003p, −log(p-value) = 8.54), *oatA* (2 for p002y, −log(p-value) = 7.75; 3 for p003p, −log(p-value) = 8.45), and *tagH* (11 for p002y, −log(p-value) = 9.47; 10 for p003p, −log(p-value) = 8.81). HsdS determines the sequence specificity of Sau1 restriction-modification (73), while OatA, or peptidoglycan O-acetyltransferase, is required for phage adsorption at least in *S. aureus* strain H (78). Prophage-associated genes (186 k-mers for phage tail fiber gene SRX477019_02350 for phage p0045, − log(p-value) = 12.21; 37 k-mers for same gene for p0040, −log(p-value) = 8.69) were the most significantly associated with two of the tested *Siphoviridae* – phage p0045 and p0040. This result agrees with the known temperate phage resistance mechanism of superinfection immunity, in which prophages express a repressor gene that prevents transcription of superinfecting phages’ lytic genes (79).

**Table 4:** GWAS summary statistics for each associated genetic element. Each value represents the number of unique genetic elements of a particular type found to be significantly associated with the phage host range phenotype.

| Phage | p0045 | p0006 | p0017 | p0017S | p002y | p003p | p0040 | pyo |
| --- | --- | --- | --- | --- | --- | --- | --- | --- |
| COGs<br>(pyseer) | 131 | 49 | 76 | 48 | 163 | 175 | 163 | 347 |
| SNPs<br>(pyseer) | 0 | 27 | 1 | 0 | 134 | 48 | 6 | 249 |
| k-mers<br>(pyseer) | 820 | 18 | 7078 | 101 | 1734 | 866 | 180 | 14 |
| SNPs<br>(treeWAS) | 0 | 1 | 0 | 0 | 0 | 1 | 1 | 4 |

TreeWAS detected 4 or fewer significant SNPs for three phages and none for phages p0045, p0017, p0017S, and p002y. Amongst significant SNPs, the majority were synonymous for each phage, with the exception of phage p0040 (**Supplemental Figure 3B**). A single nonsense mutation was detected for phage p002y. The number of significant k-mers in or near a gene detected ranged from 14 (pyo) to 7078 (p0017).

Searches using the entire set of GWAS loci for potential enriched protein-protein interactions and pathways in the STRING (64) and Gene Ontology (65) databases (using the PANTHER tool) (**Supplemental Figure 3A, Supplemental Tables S5, S6, and S7**), resulted in a biologically diverse group of functions. These included periplasmic substrate-binding (p0017S, STRING), type I restriction-modification specificity (p0017S, STRING), metal ion binding (p002y, STRING; pyo, STRING and PANTHER), ATP binding (p002y, STRING and PANTHER; pyo, STRING), amino acid metabolism (pyo, STRING and PANTHER), pyrimidine metabolism (pyo, STRING), and RNA metabolism (p0045, PANTHER). We note that the search results are limited to genes present in NCTC 8325 and must be interpreted accordingly.

### Confirmation of causal roles for novel determinants of host range

We next used molecular genetic experiments to confirm a causal role for genes discovered in the GWAS where there were no previous references in the literature for a role in *S. aureus* phage host range. The genes (*trpA* - p002y/pyseer, *phoR* - p002y, p003p, p0040/pyseer, *isdB* - p002y, p0040/pyseer, *sodM* - p002y, p003p/pyseer, *mprF/fmtC* - p002y/pyseer, and *relA* - p003p/pyseer) were selected for validation because there were available transposon mutants in the Nebraska Transposon Mutant Library (NTML) (37) and these mutants could be backcrossed into the wild-type USA300 to eliminate second site mutations. We thus could not use transposon mutants that would confer full resistance (e.g., insertions in wall teichoic biosynthesis genes *tarJ* or *tagH*) as this resistance to phage infection would prevent lysate preparation for backcrossing. Nonetheless, we backcrossed selected mutants into their isogenic background USA300 JE2 and complemented these strains with the multicopy vector pOS1-P*lgt* (66).

We assessed the USA300 JE2 background, transposon mutants, transposon mutants with empty vectors, and complemented transposon mutants for growth defects and phage resistance with the previously described high-throughput (**Figure 5** and **Supplemental Figure S6**) and efficiency of plating (EOP) assays (**Figure 5** and **Supplemental Figure S7**), respectively. No strains had growth defects respect to each other or the wild-type background (**Supplemental Figure S4**). We found significant decreases in phage resistance for all mutants in the presence of phages p0006, p0017S, p003p, and p0040 (p<0.05, Wilcoxon signed-rank test). However, when we attempted to rescue the phenotype by complementation, we only found corresponding rescue of phage resistance back towards the wild-type phenotype in *trpA*, *phoR*, *sodM*, and *fmtC* (p<0.05, Wilcoxon signed-rank test). Interestingly, the *fmtC* allele from NRS209 did not complement the *fmtC*::Tn insertion, while the *fmtC* allele from the same strain (USA300 JE2) did, suggesting allele specificity for *fmtC* in phage resistance effects. As found in growth curves (**Supplemental Figure S4**), in the high-throughput assay, for the most part, mutations and plasmids did not affect bacterial growth in the absence of phage (no phage panel in **Figure 5** and **Supplemental Figure S6**). We further evaluated bacterial survival after the high-throughput assay by measuring CFUs in assay soft agar after overnight culture for the trpA set of strains and phage p003p. As expected, surviving CFU correlated with final OD, with significantly (p<0.05, Wilcoxon signed-rank test) higher CFU and OD for USA300 JE2 than USA300 *trpA*::Tn and USA300 *trpA*::Tn pOS1 *trpA* than USA300 *trpA*::Tn pOS1, respectively (**Supplemental Figure S5**).

**Figure 5:**
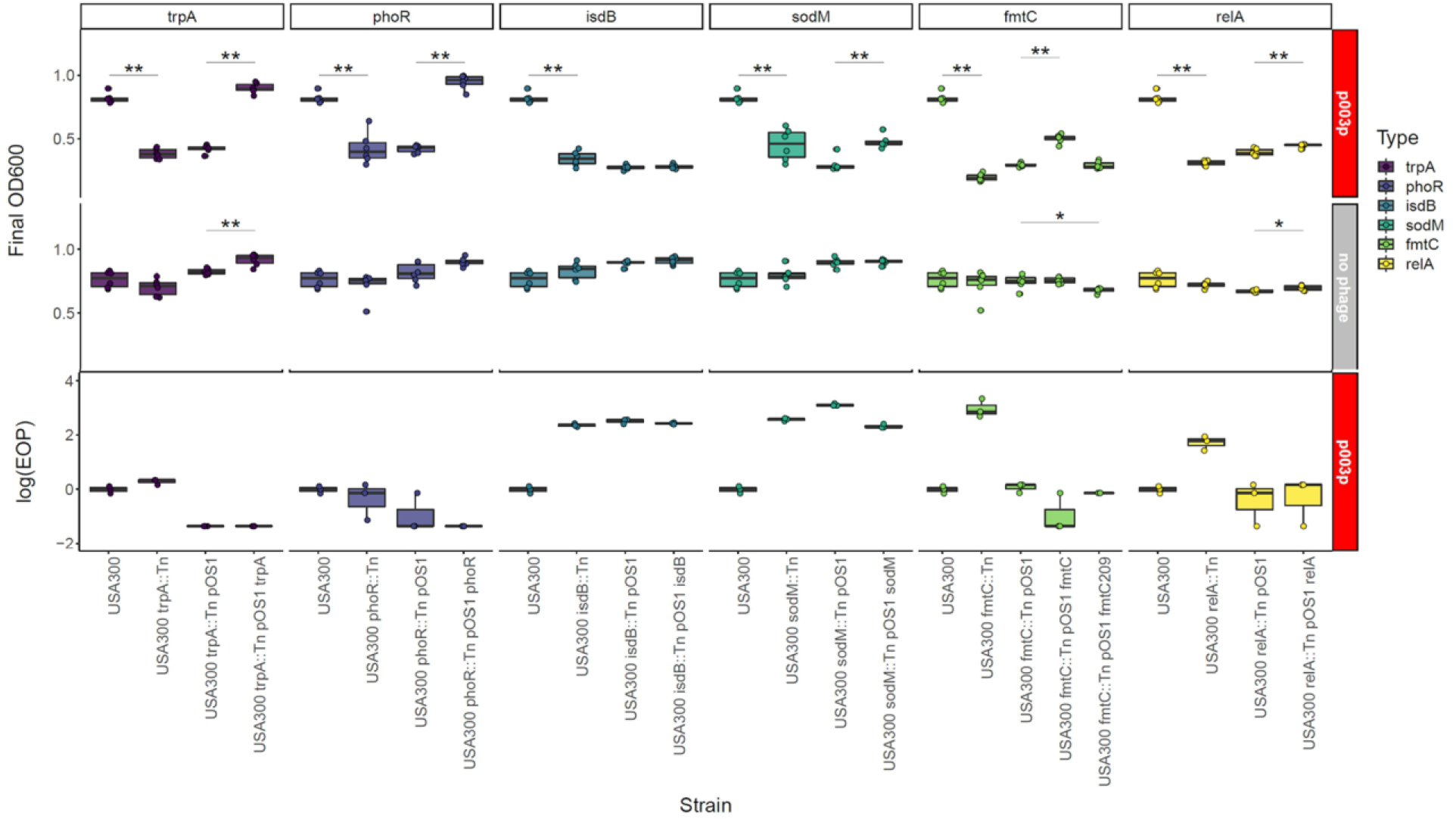
Molecular genetics validates putative phage resistance determinants. High-throughput host range assay (top) and efficiency of plating (EOP; bottom) phenotypes demonstrating genetic validation of novel GWAS phage host range determinants. Results are grouped by gene (*trpA*, *phoR*, *isdB*, *sodM*, *fmtC*, and *relA*). All assays were performed with siphovirus p003p or no phage. Each gene group includes four strains demonstrating complementation with proper controls (USA300, USA300 transposon mutant, USA300 transposon mutant with empty pOS1 vector, and USA300 transposon mutant complemented with gene in pOS1 vector). All significant (p<0.05) pairwise differences (Wilcoxon signed-rank test) are shown at the top of the corresponding boxplots.

We did not observe any significant changes in phage propagation efficiency when performing the efficiency of plating (EOP) assay on these strains, except for USA300/USA300 *trpA*::Tn pOS1 *trpA*, USA300/USA300 *phoR*::Tn pOS1, and USA300/USA300 *relA*::Tn pOS1 *relA* (p<0.05, Wilcoxon signed-rank test). EOP measures differences in plaquing, or actual infection and phage propagation. The growth-based assay measures survival despite infection. We interpreted the different results between the EOP and growth assays to indicate that these genes mostly influence survival post-infection but do not necessarily prevent infection. Taken together, these results confirmed at least six GWAS-significant genes are implicated in phage resistance for some of the eight phages but not necessarily at the level of direct interference with phage propagation.

### Host range predictive models based on significant genetic determinants explain most phenotypic variation

In order to determine the extent to which host range is predictable by the loci identified by GWAS, we constructed predictive models for qualitative host range phenotypes using random forests, gradient-boosted decision trees, and neural networks. We determined predictive accuracy for each phage host range phenotype and four different sets of predictors (presence/absence of significant genetic determinants or k-mers from GWAS result, with or without sequence type and clonal complex for corresponding strains) with 10-fold cross-validation (**Figure 6A and Supplemental Figure S8A**). In no cases were there significant differences in 10-fold cross-validation predictive accuracies between model construction methods or predictor sets used, suggesting no combination of method and predictors improved model predictive accuracy relative to another and that there is a limit to the amount of host range variation explained by the predictive models. The phages p0017S (0.83-0.87), p002y (0.81-0.88), p003p (0.83-0.92), and pyo (0.83-0.91) had the highest average predictive accuracies, followed by p0045 (0.67-0.73), p0006 (0.47-0.61), p0040 (0.42-0.61), and p0017 (0.45-0.54), respectively. We hypothesized that predictive accuracy correlated with host range distribution, expecting simpler distributions to be easier to predict and thus to have higher predictive accuracies. We thus examined the relationship between information entropy (average level of uncertainty or information in a variable’s possible outcomes) and predictive accuracy (**Figures 6B and 6C; Supplemental Figures S8B and C**). We found that predictive accuracy increased at the extremes of phenotype proportion (S, SS, R) and that information entropy was negatively correlated with predictive accuracy for all models.

**Figure 6:**
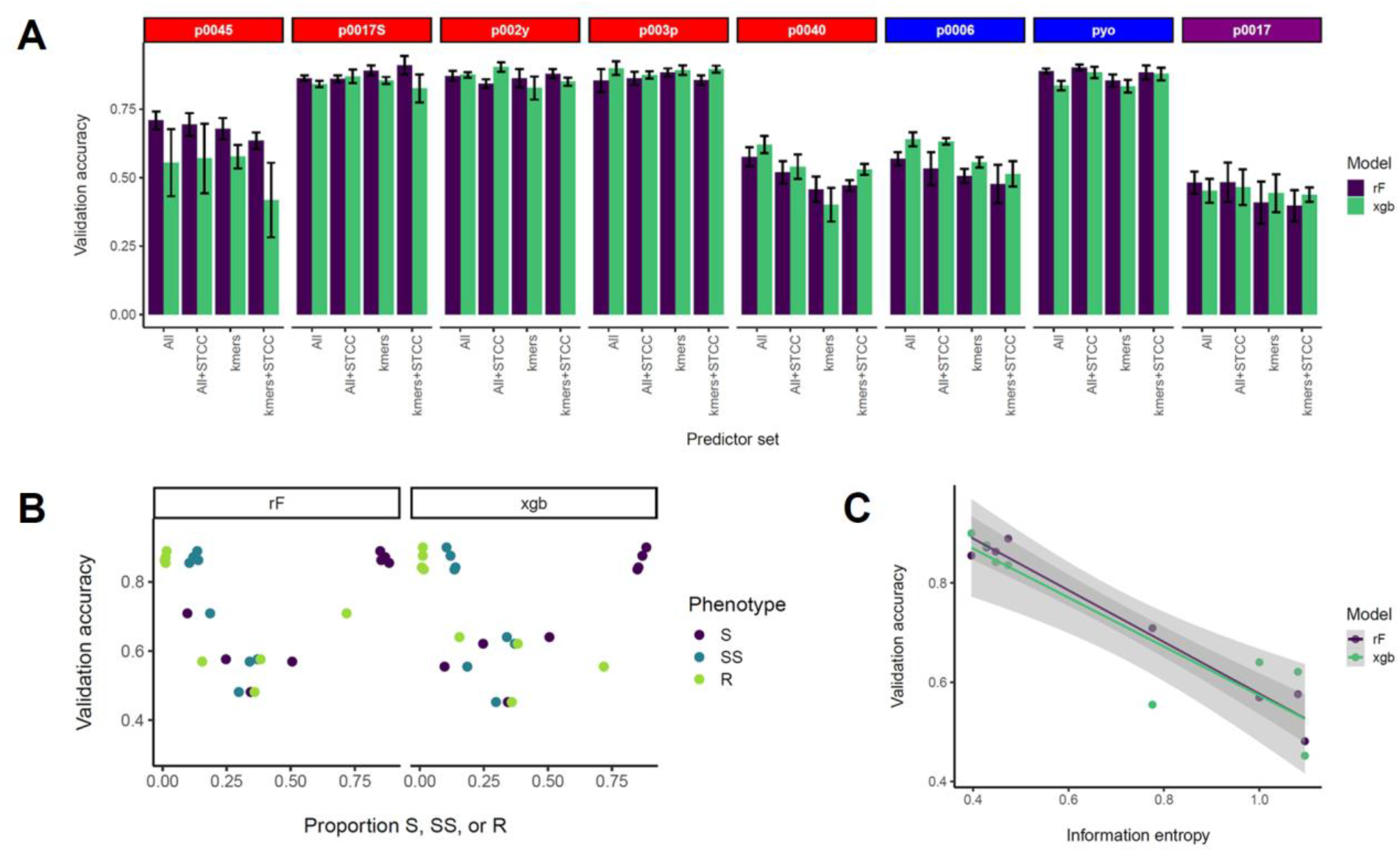
Construction of predictive models for each ternary phage resistance phenotype. Quantitative host range phenotypes were classified as S - sensitive, SS - semi-sensitive, or R - resistant based on the bins (0.1-0.4, 0.4-0.7, and 0.7 or more, respectively). *Siphoviridae* are listed in red, *Myoviridae* in blue, and *Podoviridae* in purple. A) 10-fold cross-validation predictive accuracies for each phage based on two model building methods (randomForest and XGBoost) and four sets of predictors - all significant GWAS genetic determinants (COGs, SNPs, and k-mers) for a particular phage, all determinants plus corresponding strain sequence type and clonal complex (ST and CC), significant k-mers for a particular phage, and significant k-mers plus strain ST and CC. Average accuracies of four 10-fold CV replicates are presented with one standard error above and below the mean. Validation accuracy represents the proportion of correctly identified ternary phenotypes in the validation set (one-tenth of the strain set). B) Average accuracies from four 10-fold CV replicates for each model building method and all significant GWAS determinants as predictors relative to the proportion of each ternary phenotype (S, SS, or R) amongst tested strains for the corresponding phage. Three points are shown for each validation accuracy (corresponding to each of the three possible phenotypes). C) Average accuracies from four 10-fold CV replicates for each model building method and all significant GWAS determinants as predictors relative to the information entropy for each host range phenotype, which was calculated as described in the Materials and Methods section. Information entropy was calculated with a natural logarithm in natural units (nats).

We also performed the same analyses on another predictive model statistic - the receiver operating characteristic (ROC) curve area under the curve (AUC), which measures the ability of the model to distinguish between classes (true positive and true negative). We found that gradient-boosted decision trees AUCs held uniform amongst phages, while random forest and neural network AUCs negatively correlated with information entropy (**Supplemental Figures S9 and S10**), suggesting phenotype complexity (entropy) did not affect the robustness of gradient-boosted decision tree prediction. Taken together, these results show that significant GWAS determinants from this study do not completely predict phage host range and that prediction is most effective for low complexity host range distributions, at least for random forest and neural network models.

## Discussion

Through GWAS on a diverse natural set of *S. aureus* strains we discovered numerous genetic determinants of phage host range, many of which had not been reported previously in the scientific literature. This study uses a far more diverse set of strains than the previous hypothesis-free study of *S. aureus* phage host range (36). However, our set of genetic loci still only partially explains the variation in the overall broad host ranges of our tested phages, as the predictive modeling results indicate.

We found that knockouts of six GWAS-significant genes - *trpA*, *phoR*, *isdB*, *sodM*, *fmtC*, and *relA* - increased phage sensitivity, suggesting these could be targets for phage-therapy adjunctive drugs. TrpA together with TrpB (encoding tryptophan synthase alpha and beta chains, respectively) carries out the last step in L-tryptophan biosynthesis (80). The enzymes convert indole-glycerol phosphate and serine to tryptophan and glyceraldehyde 3-phosphate (80). TrpA inactivation might then sensitize *S. aureus* to phage infection by increasing indole-glycerol phosphate levels at the expense of tryptophan. In the absence of *trpA*, built-up tryptophan biosynthesis intermediates including IGP may sensitize cells to phage infection, making *trpA* necessary for resistance. Alternatively, by reducing the total tryptophan pool, removing tryptophan biosynthesis may increase the proportion of tryptophan used to translate phage relative to host proteins, thus enhancing phage infection at the cost of host growth. Indeed, it is already known that throttling down protein synthesis with sublethal doses of ribosomal active antibiotics enhances plaque formation on MRSA lawns (81).

The PhoPR two-component system is responsible for regulating expression of phosphate uptake systems (ABC transporters) based on phosphate levels. In *S. aureus*, PhoPR is necessary for growth under phosphate-limiting conditions by regulating either phosphate transporters or other factors, depending on the environment (82). In *Bacillus subtilis*, the sensor kinase PhoR senses phosphate limitation through wall teichoic acid intermediates (83) and correspondingly represses WTA biosynthesis gene expression (84). PhoPR also upregulates glycerol-phosphate WTA degradation in *S. aureus* and *B. subtilis* to scavenge phosphate (85, 86). If all these mechanisms are present in *S. aureus*, and if there is also a pathway for degrading *S. aureus* Rbo-P WTA, PhoR activity may lead to reduced WTA under phosphate starvation, thus forming phage resistant cells. On the other hand, as for *trpA, phoR* might simply be required for properly inducing phosphate uptake necessary for survival during phage infection.

Superoxide dismutase (SodM) and phosphatidylglycerol lysyltransferase/multiple peptide resistance factor (FmtC/MprF) more likely have direct mechanistic roles in the phage infection process. SodM may be required for tolerance to cell wall stress imposed by phage infection. SodM is a Mn/Fe-dependent superoxide dismutase that converts superoxide into hydrogen peroxide and oxygen. Previous studies have shown that superoxide dismutase has affected tolerance to cell wall active antibiotics in *S. aureus* and *E. faecalis* (87, 88) and phage plaquing in *C. jejuni* (89). Superoxide dismutase transcripts were found to be upregulated upon phage infection in *E. faecalis* (90). FmtC, on the other hand, may affect the lysis step by altering cell surface charge. FmtC (MprF) alters cell surface charge first by attaching the positively charged lysine to phosphatidylglycerol through esterification with glycerol (91, 92). It then flips these modified phospholipids from the inner to the outer leaflet of the cell membrane (93). This resulting positive charge on the outer membrane confers resistance to cationic antimicrobial peptides (CAMPs) but also may alter lysis. Phage lysis depends on holin proteins which form pores in the membrane that dissipate proton-motive force and release endolysins to degrade the cell wall peptidoglycan (94–96). Because FmtC alters cell surface charge, it also could affect holin-dependent membrane depolarization, endolysin activity, or phage attachment, especially if the phage receptor-binding protein is positively charged. Interestingly enough, the *fmtC* allele from NRS209 did not complement the transposon insertion in USA300 JE2. This could indicate either a loss of function in the allele or incompatibility with some aspect of the USA300 JE2 strain.

Two of the six validated genes did not restore wild-type phenotypes upon complementation (*isdB* and *relA*). RelA, or the *relA/spoT* homolog in *S. aureus*, synthesizes (p)ppGpp in response to sensing uncharged tRNAs on the ribosome (97). Transcriptomic studies indicated *S. aureus* upregulates its *relA/spoT* homolog in response to lytic phage predation (98). RelA may contribute to phage-resistant, slow-growing cell (persister) formation (99), although studies indicate ATP depletion rather than (p)ppGpp synthesis accounts for persistence in *S. aureus* (100). IsdB, on the other hand, is part of the iron-regulated surface determinant (isd) system responsible for scavenging iron from hemoglobin (101). As experiments were conducted in rich media, the hemoglobin-iron scavenging activity of IsdB does not seem relevant, but IsdB may be an abundant surface protein, implicating it in surface occlusion. Neither of these genes are in operons, at least in USA300 JE2. It could be that the native promoters are inherently stronger than the P*lgt* promoter or are strongly upregulated during phage infection thus affecting the efficiency of complementation. We also note for all genes that there was no apparent complementation for phages p002y and pyo (**Supplemental Figure S6**). In the case of the latter two, the parental USA300 JE2 strain was already sensitive to those two phages.

These validated genes along with most other GWAS-detected host range factors were not previously reported as important in *S. aureus* phage infection, but the GWAS did identify some known factors. Such factors included WTA biosynthesis and modification genes, such as *tarP, tarJ*, and *tagH*. While TarJ and TagH are involved in WTA biosynthesis, the WTA glycosyltransferase TarP was recently shown to directly confer *Podoviridae* resistance. Capsule biosynthesis (*cap8A* and *cap8I*) (102) and peptidoglycan modification (*oatA*) genes (78) encode surface-associated functions previously implicated in *S. aureus* phage resistance. Capsule or capsule overproduction are known to confer phage resistance in *S. aureus* (7, 12), while peptidoglycan O-acetyl groups are part of the phage receptor (78). Type I restriction-modification (*hsdS*) was implicated as well, and this is a well-known mechanism for suppression of infection across clonal complexes (73). Staphylococcal pathogenicity islands (SaPIs) were not implicated most likely because these are highly specific to siphovirus helper phages, and even for possibly affected helper phages (80α), SaPI interference reduces but does not eliminate helper phage production (103). This means our high-throughput assay may not capture SaPI-level effects, as it does not directly measure phage propagation through plaquing efficiency. CRISPRs were not significant in our study either, because these are rare in *S. aureus* strains (1, 20, 21).

Our study agreed with prior work demonstrating *S. aureus* phages have broad host ranges (28–34). A major goal of our work was to prototype predictive models for host range based on genome sequence. Genome-based predictions for several antibiotic resistance phenotypes have proven to be of similar accuracy to classic laboratory-based assays (104). We found that *S. aureus* host range prediction accuracy was 40-95% depending on phage. More strains and phages will need to be added to the host range matrix to make genomic host range prediction clinically useful. The difficulty in predicting resistance may come from the large number of genes found to influence the phenotype. Resistant strains may instead have individual, unique mechanisms or other traits that simply confer phage resistance, with the exception of superinfection immunity, in which host-encoded prophages prevent infection of a cognate temperate phage by repressing its lytic genes with their *cI* repressors (79). The two phages with the highest overall resistance (p0045 and p0040 - **Figure 2**) are temperate *Siphoviridae.* Most isolated *S. aureus* strains encode prophages (105), making superinfection immunity and corresponding overall p0045 and p0040 resistance common in the tested strains.

There are limitations to performing phage host range measurement. The high-throughput assay did not measure lysis directly but also did not have the disadvantages of observer bias, low throughput, and qualitative output of the spot assay. In our host range assay, we are measuring the ability of the population overall to survive phage challenge, but this could also indicate the phage suppression of bacterial growth through some level of infection. Likewise, multiple possible sets of population dynamics confound the spot assay. Efficiency of plating (EOP), on the other hand, measures phage propagation efficiency directly by comparing phage titer on a permissive control strain to that on a test strain (69). Nonetheless, factors altering EOP still could affect any stage of the infection cycle, so EOP measurement does not suggest possible phage resistance mechanism. The ambiguity of both assays suggests examining the population dynamics of phages and identified mutants (e.g., *trpA*::Tn) during infection (i.e., adsorption rate; latent period, and burst size from one-step growth curve) would be worthy for future studies to pinpoint the specific mechanism by which that gene affects phage resistance. We also recognize that a multitude of environmental variables (temperature, multiplicity of infection, growth media) could influence the assay.

There are also some limitations inherent in GWAS approaches. Bacterial GWAS associates homoplasic variants that arise from parallel evolution or recombination with a phenotype of interest (106, 107). While bacterial GWAS can find more types of genetic events (either loss of function or gain of function; mutation, insertion, deletion, recombination, and so on, but not genes with no changes) and more broadly relevant genes and polymorphisms related to a phenotype than screening transposon mutants in a single genetic background, clonal population structure, abundant small effect variants, and genetic interactions hamper it (106). When recombination is relatively rare in a species, like *S. aureus*, large numbers of variants remain in linkage disequilibrium, making it difficult to distinguish lineage from strain-level effects. Variants linked to a causative variant may then be detected as false positives. While we have validated at least a few genes as true positives, and we expect phylogenetically hierarchical effects on host range based on reviewing past work (1), our GWAS methods also include various corrections for clonal population structure when associating variants.

Two recent studies used single gene knockout, overexpression, transcriptional suppression methods as well as global transcriptional profiling to identify phage resistance determinants in *E. coli* (108) and *Enterococcus faecalis* (90). Unlike these previous studies, our findings are not limited to one or a few genetic backgrounds, making them more widely applicable to the species and its underlying evolution. Nonetheless, extensive functional molecular genetics studies will be needed to distinguish genes that truly contribute to host range from false positives. These studies, like those in *E. coli* and *E. faecalis*, would complement the GWAS with global searches for phage resistance genes in a single genetic background, such as Tn-Seq, DUB-Seq, and CRISPR interference to identify genes required for surviving phage infection and RNA-Seq to identify genes differentially regulated in response to phage infection. Such work would both corroborate GWAS results and fill in the gaps - possible determinants not present or conserved in enough of the resistant or sensitive population.

Our results have important consequences for phage therapy, phage-small molecule combination therapy, and horizontal gene transfer in the species. Genes identified expand the set of potential combination therapies by providing additional targets to which to design small molecules to interfere with phage resistance. Already, combination phage/antibiotic therapies have shown promise for clearing biofilms and reducing emergence of antibiotic resistance in *S. aureus* (109), and ribosomal active antibiotics are known to enhance MRSA phage sensitivity at sublethal doses (81). Additionally, because the phage receptor WTA is necessary for methicillin resistance (75) and WTA inhibition resensitizes MRSA to methicillin (76), phages have the exciting possibility of inducing collateral beta-lactam sensitivity. We also cannot discount the possibility that phage resistance polymorphisms are the result of selection by other stresses besides phage infection, such as immune escape, interbacterial interactions, or antibiotic selection. Wall teichoic acid, the *S. aureus* phage receptor, for example, is also important for colonization, antibiotic resistance, and immune evasion (75, 110–115). Because we identified phage host range determinants, we also gain insights into the evolution of the *S. aureus* through horizontal gene transfer. Transduction, the transfer of host genetic material between strains by abortive phage infection, is a major mechanism of HGT (116) and recombination (117) in the species. There is a tradeoff between the need to resist phage killing and the need to adapt by gaining new virulence genes (such as Panton-Valentine leukocidin) (118) through HGT. It is possible that the most transducible strains are both more sensitive to killing by phage infection, but also more able to outcompete other strains for advantageous genetic material. The finding that even the most resistant strains (NRS148, NRS209, and NRS255) were still sensitive to two out of the eight phages may be the result of a selection for sensitivity that could be the Achilles heel of *S. aureus* when confronted by phage therapy.

## Acknowledgements

We thank Veronique Perrot and Bruce Levin for providing the pyo myophage used for host range evaluation. We also thank Bruce Levin for providing constructive comments on the manuscript. Sarah Satola and Eryn Bernardy provided VISA and CF *S. aureus* strains used for host range testing, respectively. Abraham Moller was supported by the National Science Foundation (NSF) Graduate Research Fellowship Program (GRFP). Timothy Read was supported by the National Institutes of Health (NIH) grant AI121860. Kyle Winston was supported by the Emory REAL fellowship. We thank Michelle Su and Robert Petit for assistance with GWAS methods and constructive criticism of the project.

