## Supplemental Information for "Genes influencing phage host range in *Staphylococcus aureus* on a species-wide scale"

Supplemental Figure S1: Scree plot used to pick the number of dimensions for multidimensional scaling (MDS) in pyseer COG significance analysis. The number of dimensions (PCs) picked was the least possible (1) after which the eigenvalue stabilized with respect to dimension number.

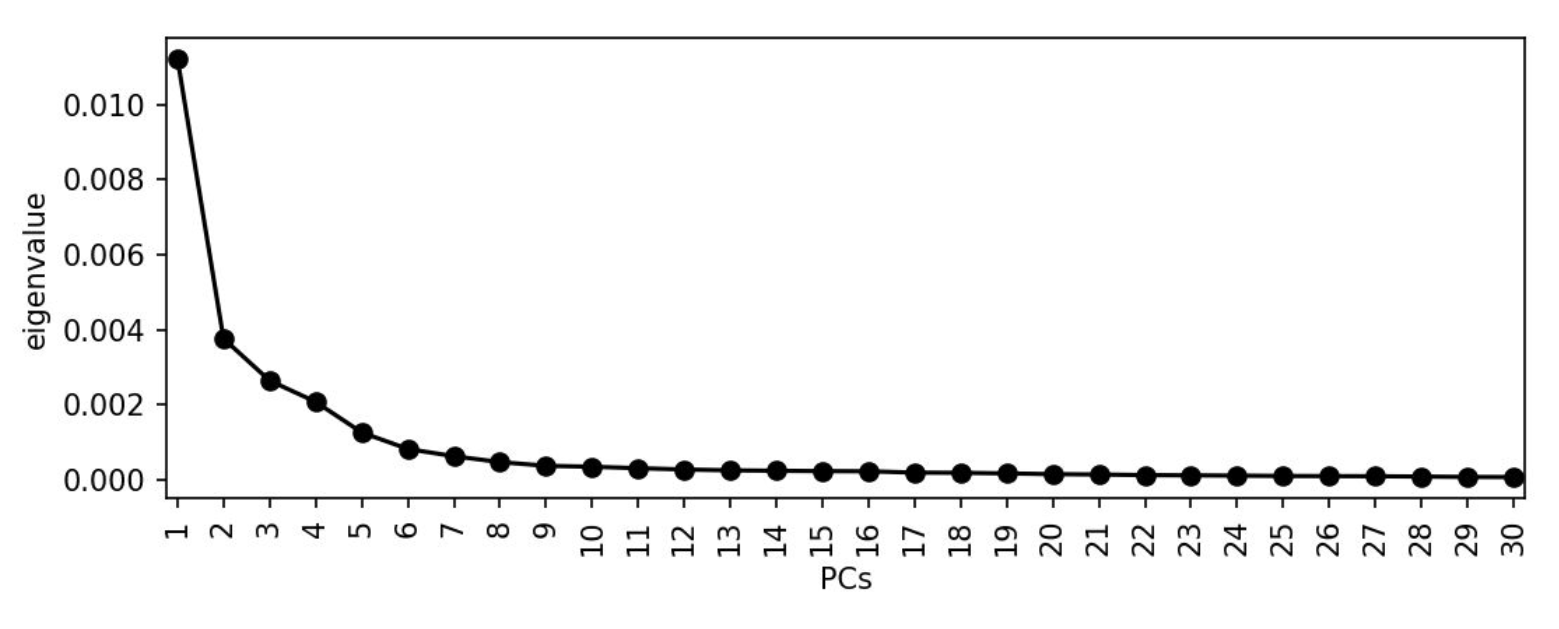

Supplemental Figure S2: Pyseer k-mer Q-Q plots for each phage (p0045, p0006, p0017, p0017S, p002y, p003p, p0040, and pyo). The observed p-values were plotted relative to the expected p-values based on the null distribution. Expected p-values were plotted with a 95% confidence interval on the diagonal. Deviation of the observed/expected curve from the diagonal indicated p-value inflation.

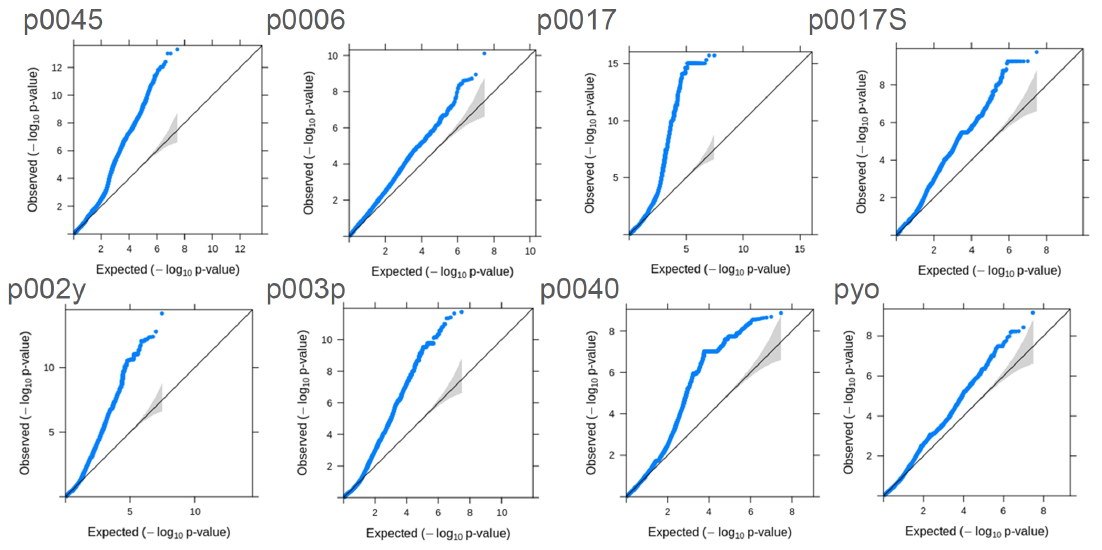

Supplemental Figure S3: GWAS approach and significant SNP annotations. A) Overview of the genome-wide association study (GWAS) workflow. Pyseer (1) associated intermediate-frequency COGs, core-genome SNPs, and k-mers with each host range phenotype, while treeWAS (2) only associated core-genome SNPs with each host range phenotype. SnpEff (3) classified mutation effects (synonymous, missense, or nonsense) from the corresponding Roary (4) gene sequence, while STRING (5) identified putative protein-protein interactions and PANTHER (6) identified enriched functions from lists of genes corresponding to each significant SNP or k-mer. B) Classification of significantly associated pyseer or treeWAS SNPs based on mutational effect (synonymous, missense, or nonsense). SnpEff annotated SNP effects based on corresponding genes identified in the tested strains’ core genome with Roary. Phage 0045 was not included as no significant SNPs were detected for its host range phenotype. *Siphoviridae* are listed in red, *Myoviridae* in blue, and *Podoviridae* in purple.

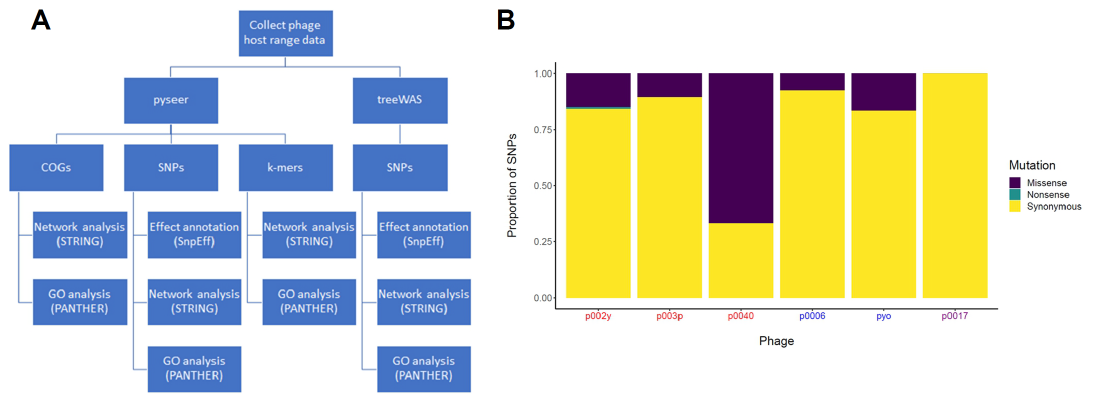

Supplemental Figure S4: Growth curves of USA300, USA300 transposon mutants (A), transposon mutants electroporated with the empty pOS1 vector (B), and transposon mutants complemented with vectors containing respective genes (C; *trpA*, *phoR*, *isdB*, *sodM*, *fmtC*, and *relA*). Strains were inoculated with a 96-pin replicator from arrayed frozen glycerol stocks into 96-well plates containing 200 µL LB/TSB 2:1 with 5 mM CaCl_2_ or the same medium supplemented with 10 μg/mL chloramphenicol in each well. We then diluted each culture 1:100 in fresh LB/TSB 2:1 with 5 mM CaCl_2_ or the same medium supplemented with 10 μg/mL chloramphenicol and collected growth curves on a BioTek Eon plate reader (37°C, 225 rpm agitation, OD_600_ measured every 10 minutes).

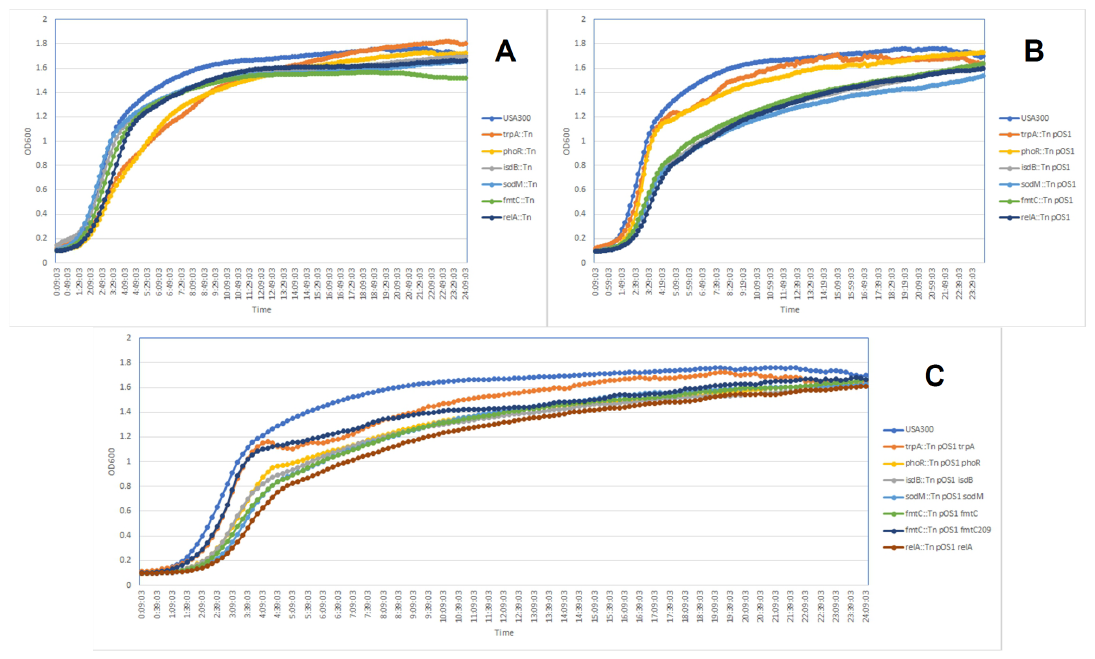

Supplemental Figure S5: Bacterial survival after completing the high-throughput host range assay (p003p against *trpA* strains). The high-throughput assay was performed for six biological replicates of USA300, USA300 *trpA*::Tn, USA300 *trpA*::Tn pOS1, and USA300 *trpA*::Tn pOS1 *trpA* strains. A) ODs were measured for the high-throughput phage host range assay replicates as described previously. B) Agar plugs were removed with toothpicks, transferred to 0.8 mL volumes of sterile TMG, and bacteria resuspended by vortexing. The resuspensions were serially diluted in TMG and 4 uL of 1e-1 through 1e-6 dilutions were spotted four times on TSA plates. Dilution plates were grown overnight at 37°C and colonies counted the following day to determine surviving CFU in each condition.

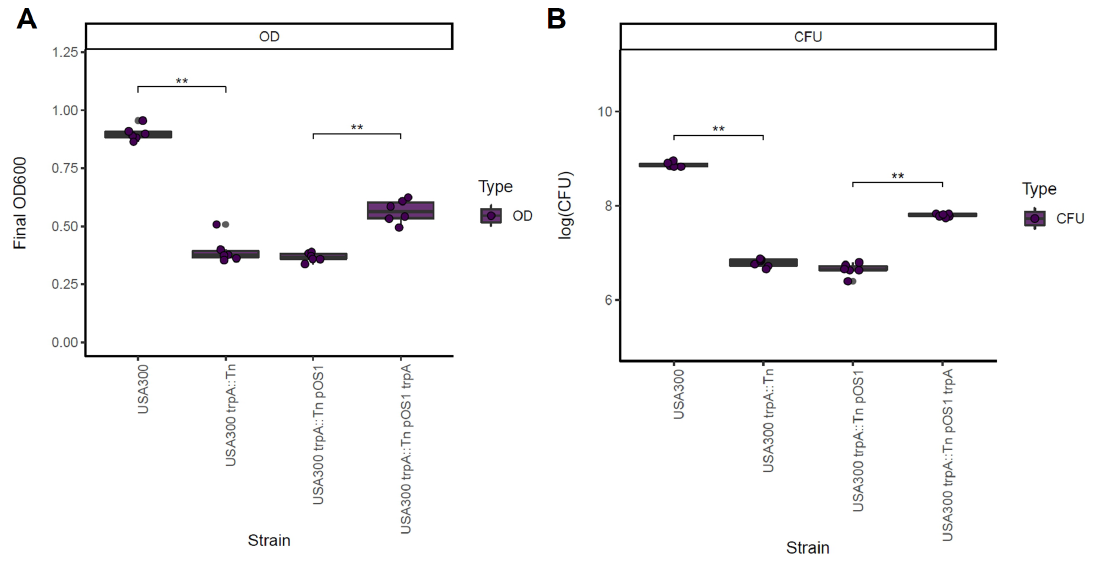

Supplemental Figure S6: High-throughput host range assay phenotypes demonstrating genetic validation of novel GWAS phage host range determinants. Results are grouped by gene (*trpA*, *phoR*, *isdB*, *sodM*, *fmtC*, and *relA*) and phage (p0045, p0017S, p003p, p0040, p0006, p002y, pyo, and no phage). Each group includes four strains demonstrating complementation with proper controls (USA300, USA300 transposon mutant, USA300 transposon mutant with empty pOS1 vector, and USA300 transposon mutant complemented with gene in pOS1 vector). All significant (p<0.05) pairwise differences (Wilcoxon signed-rank test) are shown at the top of the corresponding boxplots. *Siphoviridae* are listed in red, *Myoviridae* in blue, and the no phage control in gray.

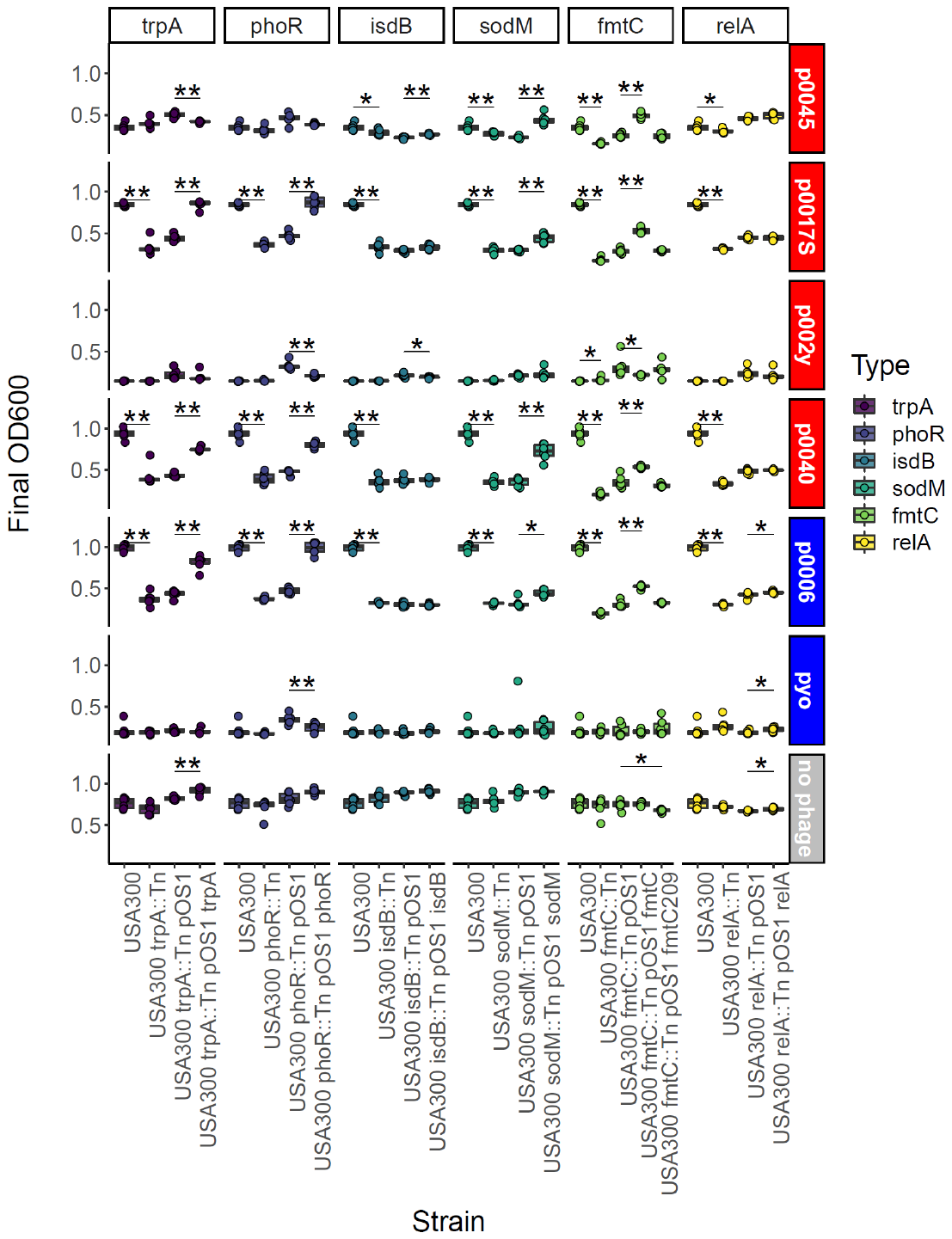

Supplemental Figure S7: Efficiency of plating (EOP) phenotypes demonstrating genetic validation of phage host range determinants. Undiluted through 1e-8 dilutions of phage were spotted (4 μL) three times on each top agar lawn, let to dry, incubated face up overnight at 37°C, and plaques counted at the lowest countable dilution. EOP was calculated relative to the average PFU/mL for the control strain, USA300 JE2. Results are grouped by gene (*trpA*, *phoR*, *isdB*, *sodM*, *fmtC*, and *relA*) and phage (p0045, p0017S, p003p, p0040, p0006, p002y, and pyo). *Siphoviridae* are listed in red and *Myoviridae* in blue. Each group includes four strains demonstrating complementation with controls (USA300, USA300 transposon mutant, USA300 transposon mutant with empty pOS1 vector, and USA300 transposon mutant complemented with gene in pOS1 vector). All significant (p<0.05) pairwise differences (Wilcoxon signed-rank test) are shown at the top of the corresponding boxplots.

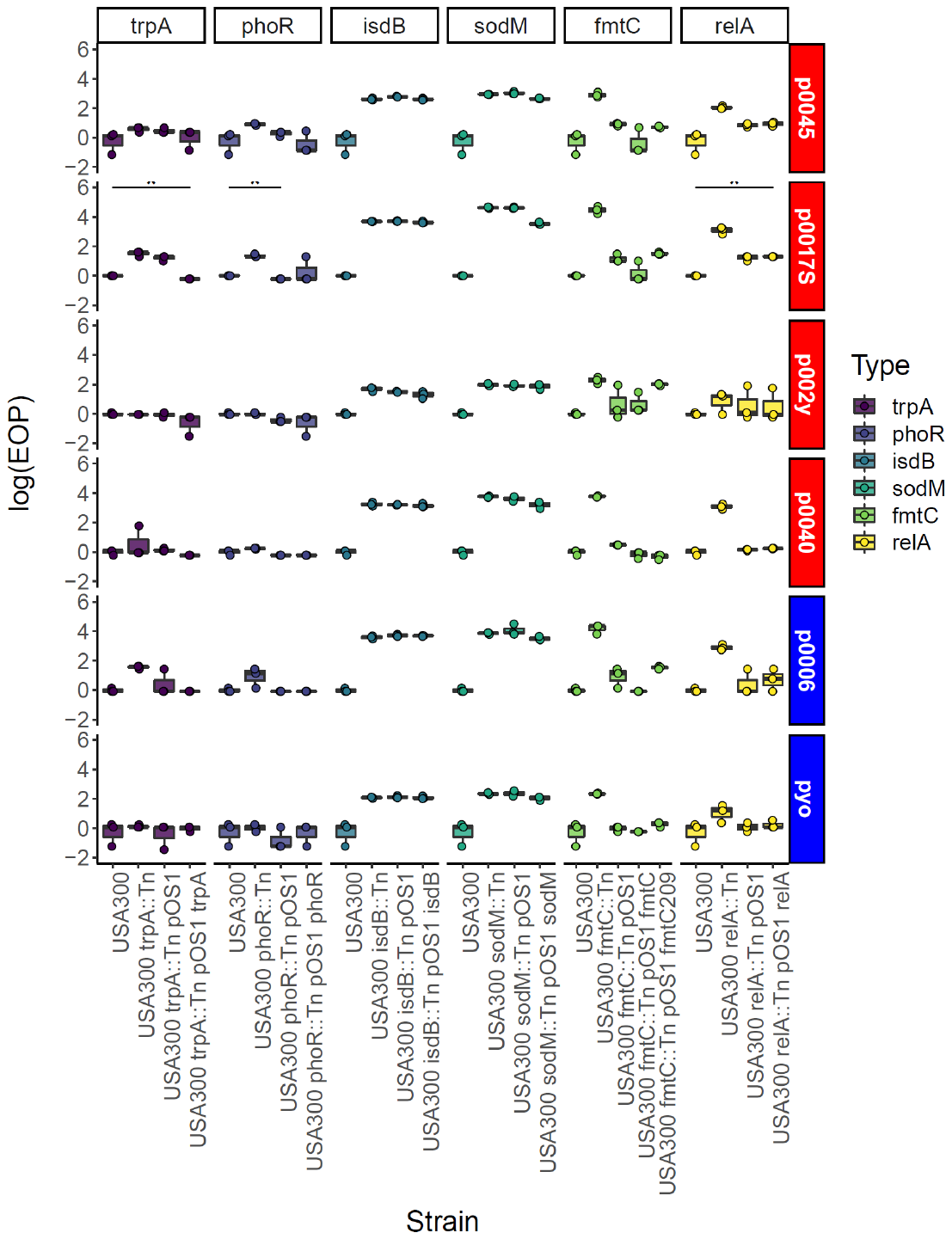

Supplemental Figure S8: Construction of neural network predictive models for each ternary phage resistance phenotype. Quantitative host range phenotypes were classified as S - sensitive, SS - semi-sensitive, or R - resistant based on the bins (0.1-0.4, 0.4-0.7, and 0.7 or more, respectively). Data preprocessing included oversampling (p0045, p0017S, p002y, p003p, or pyo), lasso regression (p0017), both (p0006), or neither (p0040). A) Predictive accuracies for each phage based on neural networks and four sets of predictors - all significant GWAS genetic determinants (COGs, SNPs, and k-mers) for a particular phage, all determinants plus corresponding strain sequence type and clonal complex (ST and CC), significant k-mers for a particular phage, and significant k-mers plus strain ST and CC. Average accuracies of four replicates are presented with one standard error above and below the mean. Validation accuracy represents the proportion of correctly identified ternary phenotypes in the validation set (30% of the strain set). B) Average accuracies from four replicates and all significant GWAS determinants as predictors relative to the proportion of each ternary phenotype (S, SS, or R) amongst tested strains for the corresponding phage. Three points on the same horizontal are shown for each validation accuracy (corresponding to each of the three possible phenotypes). C) Average accuracies from four replicates and all significant GWAS determinants as predictors relative to the information entropy for each host range phenotype, which was calculated as described in the Materials and Methods section. Information entropy was calculated with a natural logarithm in natural units (nats). *Siphoviridae* are listed in red, *Myoviridae* in blue, and *Podoviridae* in purple.

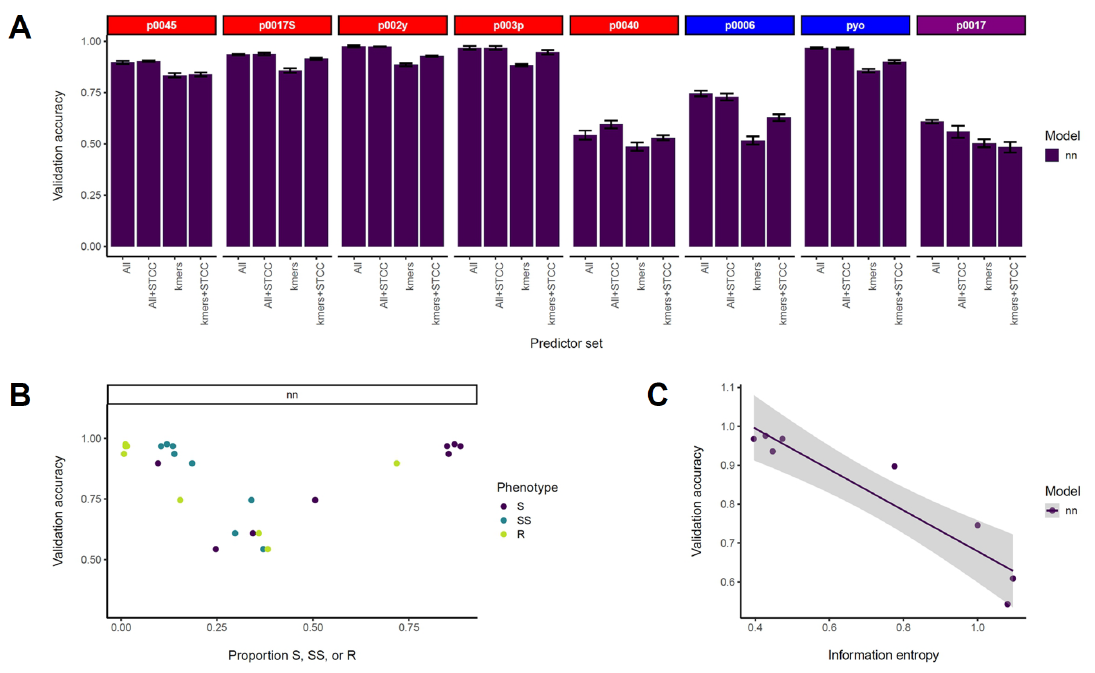

Supplemental Figure S9: Evaluation of ternary phage resistance phenotype predictive models through receiver operating characteristic-area under the curve. Quantitative host range phenotypes were classified as S - sensitive, SS - semi-sensitive, or R - resistant based on the bins 0.1-0.4, 0.4-0.7, and 0.7 or more, respectively. Data preprocessing included oversampling (p0045, p0017S, p002y, p003p, or pyo), lasso regression (p0017), both (p0006), or neither (p0040). A) 10-fold cross-validation ROC AUCs for each phage based on two model building methods (randomForest and XGBoost) and four sets of predictors - all significant GWAS genetic determinants (COGs, SNPs, and k-mers) for a particular phage, all determinants plus corresponding strain sequence type and clonal complex (ST and CC), significant k-mers for a particular phage, and significant k-mers plus strain ST and CC. Average ROC AUCs of four 10-fold CV replicates are presented with one standard error above and below the mean. B) Average ROC AUCs from four 10-fold CV replicates for each model building method and all significant GWAS determinants as predictors relative to the proportion of each ternary phenotype (S, SS, or R) amongst tested strains for the corresponding phage. Three points are shown for each ROC AUC (corresponding to each of the three possible phenotypes). C) Average ROC AUCs from four 10-fold CV replicates for each model building method and all significant GWAS determinants as predictors relative to the information entropy for each host range phenotype, which was calculated as described in the Materials and Methods section. Information entropy was calculated with a natural logarithm in natural units (nats). *Siphoviridae* are listed in red, *Myoviridae* in blue, and *Podoviridae* in purple.

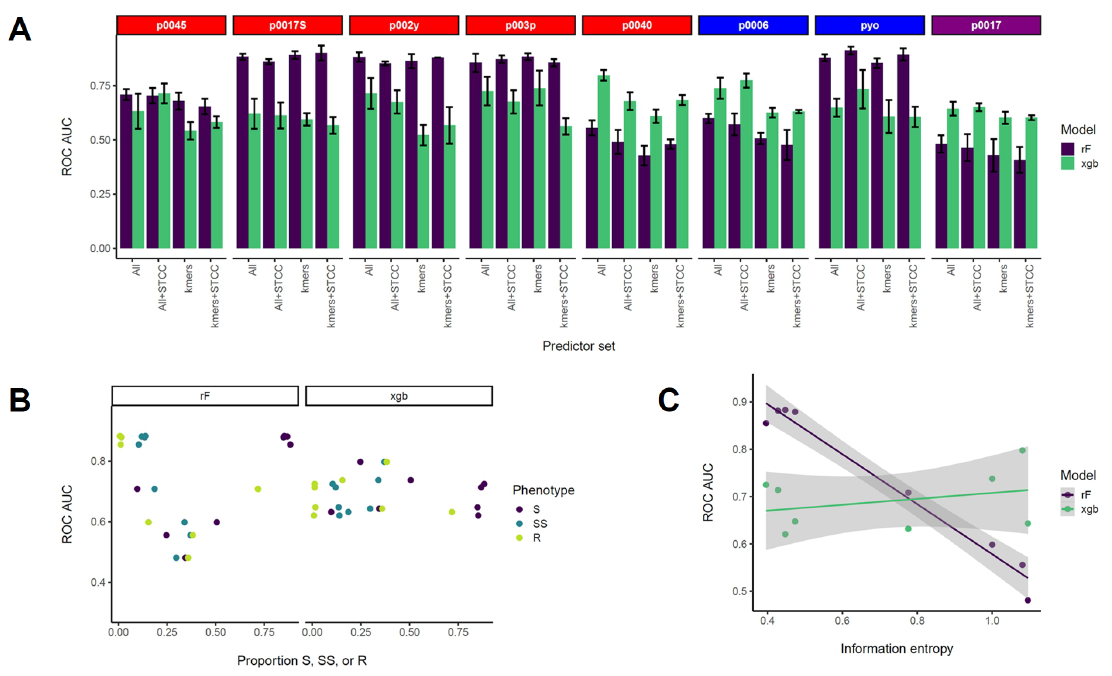

Supplemental Figure S10: Evaluation of ternary phage resistance phenotype neural network predictive models through receiver operating characteristic-area under the curve. Quantitative host range phenotypes were classified as S - sensitive, SS - semi-sensitive, or R - resistant based on the bins 0.1-0.4, 0.4-0.7, and 0.7 or more, respectively. A) ROC AUCs for each phage based on neural network models and four sets of predictors - all significant GWAS genetic determinants (COGs, SNPs, and k-mers) for a particular phage, all determinants plus corresponding strain sequence type and clonal complex (ST and CC), significant k-mers for a particular phage, and significant k-mers plus strain ST and CC. Average ROC AUCs of four replicates are presented with one standard error above and below the mean. B) Average ROC AUCs from four replicates and all significant GWAS determinants as predictors relative to the proportion of each ternary phenotype (S, SS, or R) amongst tested strains for the corresponding phage. Three points are shown for each ROC AUC (corresponding to each of the three possible phenotypes). C) Average ROC AUCs from four replicates and all significant GWAS determinants as predictors relative to the information entropy for each host range phenotype, which was calculated as described in the Materials and Methods section. Information entropy was calculated with a natural logarithm in natural units (nats). *Siphoviridae* are listed in red, *Myoviridae* in blue, and *Podoviridae* in purple.

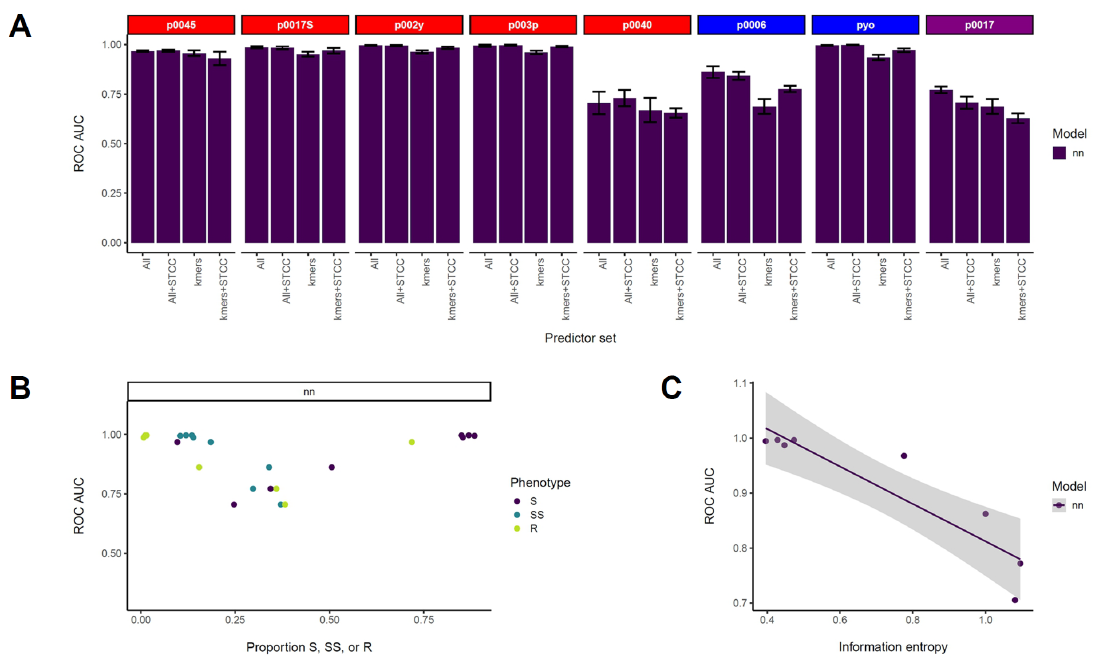

Supplemental Table S1: To determine diversity of the phages used in this study, we calculated average nucleotide identities (ANIs) with fastANI 1.31 (7). The phages were sequenced with Oxford Nanopore or Oxford Nanopore and Illumina technologies. p0017 and pyo genomes were assembled from nanopore reads with canu 2.0 (8) while p0045, p0017S, p002y, p003p, p0040, and p0006 genomes were assembled from Illumina and nanopore reads with Unicycler 0.4.8 (9).

| p0045 |  |  |  |  |  |  |  |  |
| --- | --- | --- | --- | --- | --- | --- | --- | --- |
| p0017S | 99.59 |  |  |  |  |  |  |  |
| p002y | 99.59 | 99.98 |  |  |  |  |  |  |
| p003p | 97.83 | 97.80 | 97.81 |  |  |  |  |  |
| p0040 | 99.47 | 99.85 | 99.72 | 97.75 |  |  |  |  |
| p0006 | NA | NA | NA | NA | NA |  |  |  |
| pyo (ONT) | NA | NA | NA | NA | NA | 98.88 |  |  |
| p0017 (ONT) | NA | NA | NA | NA | NA | NA | NA |  |
|  | p0045 | p0017S | p002y | p003p | p0040 | p0006 | pyo (ONT) | p0017 (ONT) |

Supplemental Table S2: Excel spreadsheet including tested *S. aureus* strain names, quantitative phenotypes, qualitative phenotypes (quantitative phenotypes from Supplemental Table S1 converted to qualitative by the OD_600_ scale 0.1-0.4 for S - sensitive, 0.4-0.7 for SS - semi-sensitive, and 0.7 or more for R - resistant), BioProject, BioSample, and SRA accessions, sequence types (STs), clonal complexes (CCs), isolation dates, and isolation locations.

Supplemental Table S3: Primers used to amplify wild-type genes corresponding to transposon mutants and clone them into pOS1-P*lgt* with splicing overlap extension (SOE)-PCR or HiFi/Gibson assembly. The annealing portion is **bolded**. Tms were calculated for NEB Q5 HF polymerase (<https://tmcalculator.neb.com/#!/batch>). *fmtC* primers were used to amplify *fmtC* from USA300 and NRS209.

| Primer | Sequence (5’ to 3’) | Tm (gene) | Tm (pOS1) |
| --- | --- | --- | --- |
| trpA LF | GGGATAAATACAATTGAGGTGAACATATGC**GCAAATGACTAAATTATTTATACC** | 55 | n/a |
| trpA LR | GGTATAAATAATTTAGTCATTTGC**GCATATGTTCACCTCAATTGTATTTATCCC** | n/a | 65 |
| trpA RF | CCAACAAACATTGAATAATTAAG**TCGAGGATCCAAACAAGGGGG** | n/a | 69 |
| trpA RR | CCCCCTTGTTTGGATCCTCGA**CTTAATTATTCAATGTTTGTTGG** | 55 | n/a |
| phoR LF | GGGATAAATACAATTGAGGTGAACATATGC**AAGAACAATGATGAAGTTTC** | 55 | n/a |
| phoR LR | GAAACTTCATCATTGTTCTT**GCATATGTTCACCTCAATTGTATTTATCCC** | n/a | 65 |
| phoR RF | CAAAGTTATTCTAAAAGATTATAAAGAATAA**TCGAGGATCCAAACAAGGGGG** | n/a | 69 |
| phoR RR | CCCCCTTGTTTGGATCCTCGA**TTATTCTTTATAATCTTTTAGAATAACTTTG** | 55 | n/a |
| isdB LF | GGGATAAATACAATTGAGGTGAACATATGC**TTTCTACAACATGAACAAAC** | 55 | n/a |
| isdB LR | GTTTGTTCATGTTGTAGAAA**GCATATGTTCACCTCAATTGTATTTATCCC** | n/a | 65 |
| isdB RF | CGTAAAAACTAATAAATCGTCT**TCGAGGATCCAAACAAGGGGG** | n/a | 69 |
| isdB RR | CCCCCTTGTTTGGATCCTCGA**AGACGATTTATTAGTTTTTACG** | 55 | n/a |
| sodM LF | GGGATAAATACAATTGAGGTGAACATATGC**GAATATACTTATGGCATTTAAATTAC** | 55 | n/a |
| sodM LR | GTAATTTAAATGCCATAAGTATATTC**GCATATGTTCACCTCAATTGTATTTATCCC** | n/a | 65 |
| sodM RF | CAAGCAGCAAAATAATATAACTTAA**TCGAGGATCCAAACAAGGGGG** | n/a | 69 |
| sodM RR | CCCCCTTGTTTGGATCCTCGA**TTAAGTTATATTATTTTGCTGCTTG** | 57 | n/a |
| fmtC LF | GGGATAAATACAATTGAGGTGAACATATGC**GTGAAAAAATGAATCAGGAAG** | 56 | n/a |
| fmtC LR | CTTCCTGATTCATTTTTTCAC**GCATATGTTCACCTCAATTGTATTTATCCC** | n/a | 65 |
| fmtC RF | CGTCACAAATAATTAAAATCC**TCGAGGATCCAAACAAGGGGG** | n/a | 69 |
| fmtC RR | CCCCCTTGTTTGGATCCTCGA**GGATTTTAATTATTTGTGACG** | 54 | n/a |
| relA LF | GGGATAAATACAATTGAGGTGAACATATGC**GTATCATATAATGAACAACGAATATCC** | 59 | n/a |
| relA LR | GGATATTCGTTGTTCATTATATGATAC**GCATATGTTCACCTCAATTGTATTTATCCC** | n/a | 65 |
| relA RF | GTTTGGAACTAGAGGTGCAAAA**TCGAGGATCCAAACAAGGGGG** | n/a | 69 |
| relA RR | CCCCCTTGTTTGGATCCTCGA**TTTTGCACCTCTAGTTCCAAAC** | 63 | n/a |

Supplemental Table S4: Significant differences (p<0.05; p-values listed in parentheses) in phage host range phenotypes between tested strains’ CCs based on Tukey HSD/one-way ANOVA tests.

| Phage | Significant comparisons |
| --- | --- |
| p0045 | CC30 vs. CC8 (3.25e-3) |
| p0006 | CC45 (1.91e-3) and 72 (6.07e-3) vs. CC1; CC45 (1.95e-3) and 72 (0.0141) vs. CC5; CC45 (4.90e-4) and CC72 (5.05e-3) vs. CC8; CC45 vs. CC30 (0.0193) |
| p0017 | CC1 (9.54e-5), 5 (2.03e-4), 30 (0.0170), and 72 (0.0140) vs. CC8 |
| p0017S | CC1 (0.0155) and 97 (0.0416) vs. CC80 |
| p002y | CC22 (0.0280), 45 (3.21e-5), and 80 (0.0177) vs. CC1; CC22 (0.0457), 45 (4.72e-6), and 80 (0.0365) vs. CC5; CC8 (2.18e-4), 30 (1.28e-3), and 97 (8.88e-3) vs. CC45 |
| p003p | CC22 (0.0142), 45 (4.20e-5), 80 (9.05e-5), 88 (0.0369), and 398 (4.91e-3) vs. CC1; CC22 (0.0227), 45 (7.53e-6), 80 (2.08e-4), and 398 (0.0112) vs. CC5; CC45 (5.12e-4), 80 (7.25e-4), and 398 (0.0431) vs. CC8; CC10 (8.55e-3), 12 (0.0148), 15 (3.39e-3), 25 (0.0483), 30 (1.01e-3), 59 (0.0106), 97 (5.43e-4), and 121 (7.92e-3) vs. CC80; CC30 (2.48e-3) and 97 (0.0102) vs. CC45; CC398 (0.0343) vs. CC97 |
| p0040 | CC5 (5.83e-5), 30 (1.95e-6), and 121 (9.04e-5) vs. CC8 |
| pyo | CC45 (2.96e-3), 80 (3.86e-3), 88 (8.90e-3), 121 (8.94e-3), and 398 (0.0167) vs. CC1; CC30 (0.0138), 45 (3.10e-5), 80 (2.47e-3), 88 (6.01e-3), 121 (3.54e-5), and 398 (7.79e-3) vs. CC5; CC45 (1.19e-3), 80 (6.58e-3), 88 (0.0150), 121 (2.58e-3), and 398 (0.0275) vs. CC8; CC22 (0.0482) and 30 (0.0492) vs. CC80 |

#### Supplemental Table S5: Summary statistics for protein-protein interaction networks identified with STRING amongst genes corresponding to significant SNPs or k-mers (inside or adjacent to genes). PPI enrichment p-value corresponds to the likelihood nodes and edges would be selected from the *S. aureus* database by chance.

| Phage | p0045 | p0006 | p0017 | p0017S | p002y | p003p | p0040 | pyo |
| --- | --- | --- | --- | --- | --- | --- | --- | --- |
| number of nodes | 39 | 33 | 49 | 25 | 405 | 214 | 25 | 164 |
| number of edges | 18 | 6 | 26 | 6 | 1075 | 347 | 10 | 304 |
| average node degree | 0.92 | 0.36 | 1.06 | 0.48 | 5.31 | 3.24 | 0.8 | 3.71 |
| avg. local clustering coefficient | 0.36 | 0.30 | 0.33 | 0.21 | 0.38 | 0.38 | 0.52 | 0.38 |
| expected number of edges | 6 | 6 | 18 | 4 | 930 | 287 | 5 | 243 |
| PPI enrichment p-value | 0.00015 | 0.62 | 0.038 | 0.23 | 1.77E-06 | 3.24E-04 | 0.036 | 9.45E-05 |

Supplemental Table S6: Functions enriched amongst gene sets analyzed with STRING databases. Sets of gene names corresponding to significant COGs, SNPs, and k-mers were created as described in the Materials and Methods section.

| Term ID | term description | observed gene count | background gene count | false discovery rate |
| --- | --- | --- | --- | --- |
| **p0045** |  |  |  |  |
| PFAM Protein Domains | | |  |  |
| PF10651 | Domain of unknown function (DUF2479) | 2 | 2 | 0.041 |
| **p0006** |  |  |  |  |
| none |  |  |  |  |
| **p0017** |  |  |  |  |
| SMART Protein Domains | | |  |  |
| SM00062 | Bacterial periplasmic substrate-binding proteins | 2 | 2 | 0.015 |
| SM00287 | Bacterial SH3 domain homologues | 2 | 4 | 0.0183 |
| Reference publications | | |  |  |
| PMID:29270158 | (2017) Commercial Biocides Induce Transfer of Prophage Phi13 from Human Strains of Staphylococcus aureus to Livestock CC398. | 4 | 6 | 0.007 |
| PMID:28515479 | (2017) Acquisition of virulence factors in livestock-associated MRSA: Lysogenic conversion of CC398 strains by virulence gene-containing phages. | 3 | 4 | 0.0397 |
| **p0017S** |  |  |  |  |
| PFAM Protein Domains | | |  |  |
| PF01420 | Type I restriction modification DNA specificity domain | 2 | 2 | 0.0149 |
| INTERPRO Protein Domains and Features | | | |  |
| IPR000055 | Restriction endonuclease, type I, HsdS | 2 | 2 | 0.02 |
| **p002y** |  |  |  |  |
| Biological Process (GO) | | |  |  |
| GO:0008152 | metabolic process | 105 | 530 | 0.0464 |
| Molecular Function (GO) | | |  |  |
| GO:0046872 | metal ion binding | 36 | 135 | 0.0403 |
| GO:0005488 | binding | 81 | 393 | 0.0403 |
| GO:0003824 | catalytic activity | 91 | 451 | 0.0403 |
| GO:0043167 | ion binding | 58 | 270 | 0.0425 |
| UniProt Keywords | |  |  |  |
| KW-0067 | ATP-binding | 56 | 244 | 0.0442 |
| **p003p** |  |  |  |  |
| none |  |  |  |  |
| **p0040** |  |  |  |  |
| KEGG Pathways | |  |  |  |
| sauw00051 | Fructose and mannose metabolism | 3 | 15 | 0.0072 |
| PFAM Protein Domains | | |  |  |
| PF10651 | Domain of unknown function (DUF2479) | 2 | 2 | 0.0094 |
| PF05031 | Iron Transport-associated domain | 2 | 4 | 0.0116 |
| INTERPRO Protein Domains and Features | | | |  |
| IPR018913 | BppU, N-terminal | 2 | 2 | 0.0208 |
| IPR037250 | NEAT domain superfamily | 2 | 4 | 0.0257 |
| IPR006635 | NEAT domain | 2 | 4 | 0.0257 |
| SMART Protein Domains | | |  |  |
| SM00725 | NEAr Transporter domain | 2 | 4 | 0.0107 |
| **pyo** |  |  |  |  |
| Keyword |  |  |  |  |
| KW-0479 | Metal-binding | 33 | 218 | 3.66E-05 |
| KW-0963 | Cytoplasm | 38 | 310 | 0.00025 |
| KW-0560 | Oxidoreductase | 20 | 133 | 0.004 |
| KW-0460 | Magnesium | 14 | 83 | 0.0131 |
| KW-0808 | Transferase | 29 | 279 | 0.0243 |
| KW-0143 | Chaperone | 6 | 19 | 0.0311 |
| KW-0456 | Lyase | 12 | 74 | 0.0311 |
| KEGG |  |  |  |  |
| sauw01100 | Metabolic pathways | 57 | 424 | 1.00E-08 |
| sauw01110 | Biosynthesis of secondary metabolites | 29 | 214 | 0.00041 |
| sauw00240 | Pyrimidine metabolism | 9 | 40 | 0.0174 |
| sauw01130 | Biosynthesis of antibiotics | 20 | 164 | 0.0185 |
| sauw00130 | Ubiquinone and other terpenoid-quinone biosynthesis | 4 | 8 | 0.0263 |
| sauw00260 | Glycine, serine and threonine metabolism | 7 | 29 | 0.0263 |
| sauw01120 | Microbial metabolism in diverse environments | 16 | 123 | 0.0263 |
| sauw03070 | Bacterial secretion system | 4 | 9 | 0.03 |
| Component | |  |  |  |
| GO:0005737 | cytoplasm | 41 | 334 | 3.23E-05 |
| GO:0044424 | intracellular part | 43 | 351 | 3.23E-05 |
| GO:0044464 | cell part | 49 | 495 | 0.00034 |
| GO:0044444 | cytoplasmic part | 16 | 128 | 0.0216 |
| Function |  |  |  |  |
| GO:0043167 | ion binding | 39 | 270 | 1.05E-05 |
| GO:0046872 | metal ion binding | 26 | 135 | 1.05E-05 |
| GO:0003824 | catalytic activity | 51 | 451 | 2.17E-05 |
| GO:0005488 | binding | 45 | 393 | 7.96E-05 |
| GO:0036094 | small molecule binding | 25 | 191 | 0.0029 |
| GO:0097159 | organic cyclic compound binding | 33 | 305 | 0.0052 |
| GO:1901363 | heterocyclic compound binding | 33 | 305 | 0.0052 |
| GO:0046914 | transition metal ion binding | 8 | 26 | 0.0056 |
| GO:0016740 | transferase activity | 20 | 147 | 0.006 |
| GO:0048037 | cofactor binding | 11 | 52 | 0.006 |
| GO:0043168 | anion binding | 22 | 179 | 0.0089 |
| GO:0000287 | magnesium ion binding | 5 | 10 | 0.0107 |
| GO:0005515 | protein binding | 6 | 17 | 0.0123 |
| GO:0000166 | nucleotide binding | 21 | 177 | 0.0146 |
| GO:0008144 | drug binding | 17 | 147 | 0.0471 |
| GO:0097367 | carbohydrate derivative binding | 18 | 159 | 0.0471 |
| GO:0005524 | ATP binding | 16 | 136 | 0.0495 |
| Process |  |  |  |  |
| GO:0009987 | cellular process | 58 | 519 | 3.41E-05 |
| GO:0008152 | metabolic process | 58 | 530 | 3.45E-05 |
| GO:0044237 | cellular metabolic process | 53 | 469 | 3.45E-05 |
| GO:0044281 | small molecule metabolic process | 32 | 208 | 3.45E-05 |
| GO:0071704 | organic substance metabolic process | 55 | 492 | 3.45E-05 |
| GO:0006807 | nitrogen compound metabolic process | 48 | 423 | 8.62E-05 |
| GO:0044238 | primary metabolic process | 49 | 439 | 8.62E-05 |
| GO:1901564 | organonitrogen compound metabolic process | 36 | 306 | 0.0011 |
| GO:0034641 | cellular nitrogen compound metabolic process | 35 | 316 | 0.0041 |
| GO:0006082 | organic acid metabolic process | 20 | 135 | 0.0054 |
| GO:0046483 | heterocycle metabolic process | 29 | 248 | 0.006 |
| GO:1901576 | organic substance biosynthetic process | 35 | 326 | 0.006 |
| GO:0019752 | carboxylic acid metabolic process | 18 | 118 | 0.0063 |
| GO:1901565 | organonitrogen compound catabolic process | 8 | 25 | 0.0065 |
| GO:0006725 | cellular aromatic compound metabolic process | 28 | 243 | 0.0067 |
| GO:0044249 | cellular biosynthetic process | 34 | 322 | 0.0067 |
| GO:1901360 | organic cyclic compound metabolic process | 29 | 256 | 0.0067 |
| GO:0044282 | small molecule catabolic process | 9 | 34 | 0.0069 |
| GO:0009056 | catabolic process | 12 | 61 | 0.0072 |
| GO:0044248 | cellular catabolic process | 11 | 52 | 0.0072 |
| GO:0016054 | organic acid catabolic process | 7 | 22 | 0.0118 |
| GO:0008610 | lipid biosynthetic process | 8 | 30 | 0.0122 |
| GO:1901135 | carbohydrate derivative metabolic process | 16 | 109 | 0.0122 |
| GO:0006139 | nucleobase-containing compound metabolic process | 24 | 208 | 0.0131 |
| GO:0009063 | cellular amino acid catabolic process | 6 | 16 | 0.0131 |
| GO:1901606 | alpha-amino acid catabolic process | 6 | 16 | 0.0131 |
| GO:0055086 | nucleobase-containing small molecule metabolic process | 12 | 69 | 0.014 |
| GO:1901575 | organic substance catabolic process | 11 | 60 | 0.0153 |
| GO:0044255 | cellular lipid metabolic process | 8 | 33 | 0.0162 |
| GO:0006520 | cellular amino acid metabolic process | 13 | 83 | 0.0184 |
| GO:0006796 | phosphate-containing compound metabolic process | 15 | 110 | 0.0268 |
| GO:1901605 | alpha-amino acid metabolic process | 10 | 58 | 0.0319 |
| GO:0019637 | organophosphate metabolic process | 12 | 83 | 0.0438 |
| GO:0006457 | protein folding | 4 | 9 | 0.0454 |

Supplemental Table S7: Functions enriched amongst gene sets analyzed with PANTHER databases. Sets of gene names corresponding to significant COGs, SNPs, and k-mers were created as described in the Materials and Methods section. FDR means False Discovery Rate.

| PANTHER GO-Slim Biological Process | Staphylococcus aureus - REFLIST (2889) | p0045 (24) | p0045 (expected) | p0045 (over/under) | p0045 (fold Enrichment) | p0045 (raw P-value) | p0045 (FDR) |
| --- | --- | --- | --- | --- | --- | --- | --- |
| regulation of nucleic acid-templated transcription (GO:1903506) | 45 | 4 | 0.37 | + | 10.7 | 5.87E-04 | 1.17E-01 |
| regulation of RNA biosynthetic process (GO:2001141) | 45 | 4 | 0.37 | + | 10.7 | 5.87E-04 | 8.74E-02 |
| nucleic acid-templated transcription (GO:0097659) | 46 | 4 | 0.38 | + | 10.47 | 6.34E-04 | 7.56E-02 |
| regulation of RNA metabolic process (GO:0051252) | 46 | 4 | 0.38 | + | 10.47 | 6.34E-04 | 6.30E-02 |
| regulation of nucleobase-containing compound metabolic process (GO:0019219) | 47 | 4 | 0.39 | + | 10.24 | 6.84E-04 | 5.83E-02 |
| RNA biosynthetic process (GO:0032774) | 47 | 4 | 0.39 | + | 10.24 | 6.84E-04 | 5.10E-02 |
| regulation of macromolecule biosynthetic process (GO:0010556) | 51 | 4 | 0.42 | + | 9.44 | 9.13E-04 | 6.05E-02 |
| regulation of biosynthetic process (GO:0009889) | 51 | 4 | 0.42 | + | 9.44 | 9.13E-04 | 5.44E-02 |
| regulation of cellular biosynthetic process (GO:0031326) | 51 | 4 | 0.42 | + | 9.44 | 9.13E-04 | 4.95E-02 |
| regulation of cellular macromolecule biosynthetic process (GO:2000112) | 51 | 4 | 0.42 | + | 9.44 | 9.13E-04 | 4.54E-02 |
| regulation of nitrogen compound metabolic process (GO:0051171) | 52 | 4 | 0.43 | + | 9.26 | 9.78E-04 | 4.48E-02 |
| regulation of primary metabolic process (GO:0080090) | 52 | 4 | 0.43 | + | 9.26 | 9.78E-04 | 4.16E-02 |
| regulation of gene expression (GO:0010468) | 52 | 4 | 0.43 | + | 9.26 | 9.78E-04 | 3.89E-02 |
| regulation of cellular metabolic process (GO:0031323) | 53 | 4 | 0.44 | + | 9.08 | 1.05E-03 | 3.90E-02 |
| regulation of macromolecule metabolic process (GO:0060255) | 53 | 4 | 0.44 | + | 9.08 | 1.05E-03 | 3.67E-02 |
| regulation of metabolic process (GO:0019222) | 54 | 4 | 0.45 | + | 8.92 | 1.12E-03 | 3.70E-02 |
| macromolecule biosynthetic process (GO:0009059) | 123 | 6 | 1.02 | + | 5.87 | 4.68E-04 | 2.79E-01 |
| cellular macromolecule biosynthetic process (GO:0034645) | 123 | 6 | 1.02 | + | 5.87 | 4.68E-04 | 1.39E-01 |
| RNA metabolic process (GO:0016070) | 104 | 5 | 0.86 | + | 5.79 | 1.61E-03 | 4.57E-02 |
| nucleic acid metabolic process (GO:0090304) | 148 | 6 | 1.23 | + | 4.88 | 1.20E-03 | 3.78E-02 |
| Nucleobase-  containing compound metabolic process (GO:0006139) | 211 | 7 | 1.75 | + | 3.99 | 1.37E-03 | 4.09E-02 |
| PANTHER Protein Class | Staphylococcus aureus - REFLIST (2889) | p0017 (38) | p0017 (expected) | p0017 (over/under) | p0017 (fold Enrichment) | p0017 (raw P-value) | p0017 (FDR) |
| protein class (PC00000) | 986 | 25 | 12.97 | + | 1.93 | 9.62E-05 | 4.95E-03 |
| Unclassified (UNCLASSIFIED) | 1903 | 13 | 25.03 | - | 0.52 | 9.62E-05 | 9.91E-03 |
| GO biological process complete | Staphylococcus aureus - REFLIST (2889) | p002y (326) | p002y (expected) | p002y (over/under) | p002y (fold Enrichment) | p002y (raw P-value) | p002y (FDR) |
| organic substance metabolic process (GO:0071704) | 878 | 143 | 99.08 | + | 1.44 | 1.62E-06 | 6.35E-04 |
| cellular metabolic process (GO:0044237) | 837 | 136 | 94.45 | + | 1.44 | 4.19E-06 | 1.31E-03 |
| metabolic process (GO:0008152) | 1030 | 167 | 116.23 | + | 1.44 | 6.30E-08 | 3.29E-05 |
| primary metabolic process (GO:0044238) | 748 | 121 | 84.41 | + | 1.43 | 3.14E-05 | 7.01E-03 |
| nitrogen compound metabolic process (GO:0006807) | 717 | 115 | 80.91 | + | 1.42 | 7.77E-05 | 1.52E-02 |
| biological_process (GO:0008150) | 1496 | 237 | 168.81 | + | 1.4 | 3.16E-13 | 2.48E-10 |
| cellular process (GO:0009987) | 947 | 146 | 106.86 | + | 1.37 | 2.00E-05 | 5.21E-03 |
| Unclassified (UNCLASSIFIED) | 1393 | 89 | 157.19 | - | 0.57 | 3.16E-13 | 4.95E-10 |
| GO molecular function complete | Staphylococcus aureus - REFLIST (2889) | p002y (326) | p002y (expected) | p002y (over/under) | p002y (fold Enrichment) | p002y (raw P-value) | p002y (FDR) |
| ATP binding (GO:0005524) | 275 | 58 | 31.03 | + | 1.87 | 1.37E-05 | 1.23E-03 |
| adenyl ribonucleotide binding (GO:0032559) | 276 | 58 | 31.14 | + | 1.86 | 2.01E-05 | 1.38E-03 |
| adenyl nucleotide binding (GO:0030554) | 277 | 58 | 31.26 | + | 1.86 | 2.07E-05 | 1.34E-03 |
| DNA binding (GO:0003677) | 215 | 45 | 24.26 | + | 1.85 | 2.28E-04 | 1.21E-02 |
| carbohydrate derivative binding (GO:0097367) | 320 | 66 | 36.11 | + | 1.83 | 5.86E-06 | 8.54E-04 |
| drug binding (GO:0008144) | 311 | 64 | 35.09 | + | 1.82 | 1.01E-05 | 1.18E-03 |
| purine ribonucleoside triphosphate binding (GO:0035639) | 302 | 62 | 34.08 | + | 1.82 | 1.80E-05 | 1.50E-03 |
| ribonucleotide binding (GO:0032553) | 312 | 64 | 35.21 | + | 1.82 | 1.06E-05 | 1.12E-03 |
| purine ribonucleotide binding (GO:0032555) | 303 | 62 | 34.19 | + | 1.81 | 1.85E-05 | 1.44E-03 |
| purine nucleotide binding (GO:0017076) | 304 | 62 | 34.3 | + | 1.81 | 1.92E-05 | 1.40E-03 |
| anion binding (GO:0043168) | 378 | 74 | 42.65 | + | 1.73 | 7.40E-06 | 9.58E-04 |
| nucleotide binding (GO:0000166) | 382 | 72 | 43.11 | + | 1.67 | 3.55E-05 | 2.18E-03 |
| nucleoside phosphate binding (GO:1901265) | 382 | 72 | 43.11 | + | 1.67 | 3.55E-05 | 2.07E-03 |
| ion binding (GO:0043167) | 569 | 107 | 64.21 | + | 1.67 | 1.37E-07 | 2.29E-05 |
| small molecule binding (GO:0036094) | 427 | 80 | 48.18 | + | 1.66 | 1.33E-05 | 1.29E-03 |
| nucleic acid binding (GO:0003676) | 353 | 66 | 39.83 | + | 1.66 | 1.20E-04 | 6.65E-03 |
| hydrolase activity (GO:0016787) | 364 | 65 | 41.07 | + | 1.58 | 4.08E-04 | 2.07E-02 |
| heterocyclic compound binding (GO:1901363) | 710 | 126 | 80.12 | + | 1.57 | 1.19E-07 | 2.77E-05 |
| organic cyclic compound binding (GO:0097159) | 710 | 126 | 80.12 | + | 1.57 | 1.19E-07 | 2.31E-05 |
| binding (GO:0005488) | 898 | 156 | 101.33 | + | 1.54 | 2.60E-09 | 7.57E-07 |
| catalytic activity (GO:0003824) | 1105 | 189 | 124.69 | + | 1.52 | 1.10E-11 | 4.27E-09 |
| molecular_function (GO:0003674) | 1567 | 256 | 176.82 | + | 1.45 | 6.25E-18 | 3.64E-15 |
| Unclassified (UNCLASSIFIED) | 1322 | 70 | 149.18 | - | 0.47 | 6.25E-18 | 7.28E-15 |
| PANTHER GO-Slim Molecular Function | Staphylococcus aureus - REFLIST (2889) | p002y (326) | p002y (expected) | p002y (over/under) | p002y (fold Enrichment) | p002y (raw P-value) | p002y (FDR) |
| catalytic activity (GO:0003824) | 498 | 90 | 56.2 | + | 1.6 | 1.44E-05 | 1.18E-03 |
| molecular_function (GO:0003674) | 729 | 122 | 82.26 | + | 1.48 | 4.48E-06 | 5.51E-04 |
| Unclassified (UNCLASSIFIED) | 2160 | 204 | 243.74 | - | 0.84 | 4.48E-06 | 1.10E-03 |
| PANTHER Protein Class | Staphylococcus aureus - REFLIST (2889) | p002y (326) | p002y (expected) | p002y (over/under) | p002y (fold Enrichment) | p002y (raw P-value) | p002y (FDR) |
| protein class (PC00000) | 986 | 172 | 111.26 | + | 1.55 | 7.79E-11 | 4.01E-09 |
| metabolite interconversion enzyme (PC00262) | 473 | 78 | 53.37 | + | 1.46 | 1.08E-03 | 3.69E-02 |
| Unclassified (UNCLASSIFIED) | 1903 | 154 | 214.74 | - | 0.72 | 7.79E-11 | 8.02E-09 |
| GO biological process complete | Staphylococcus aureus - REFLIST (2889) | p003p (181) | p003p (expected) | p003p (over/under) | p003p (fold Enrichment) | p003p (raw P-value) | p003p (FDR) |
| Nucleobase-  containing compound metabolic process (GO:0006139) | 331 | 42 | 20.74 | + | 2.03 | 1.74E-05 | 5.45E-03 |
| cellular aromatic compound metabolic process (GO:0006725) | 394 | 46 | 24.68 | + | 1.86 | 4.34E-05 | 1.13E-02 |
| cellular nitrogen compound metabolic process (GO:0034641) | 491 | 56 | 30.76 | + | 1.82 | 8.37E-06 | 4.36E-03 |
| heterocycle metabolic process (GO:0046483) | 408 | 46 | 25.56 | + | 1.8 | 1.30E-04 | 2.26E-02 |
| organic cyclic compound metabolic process (GO:1901360) | 418 | 46 | 26.19 | + | 1.76 | 2.34E-04 | 3.33E-02 |
| nitrogen compound metabolic process (GO:0006807) | 717 | 69 | 44.92 | + | 1.54 | 1.43E-04 | 2.24E-02 |
| cellular metabolic process (GO:0044237) | 837 | 78 | 52.44 | + | 1.49 | 1.05E-04 | 2.05E-02 |
| metabolic process (GO:0008152) | 1030 | 95 | 64.53 | + | 1.47 | 1.03E-05 | 4.05E-03 |
| cellular process (GO:0009987) | 947 | 86 | 59.33 | + | 1.45 | 8.91E-05 | 1.99E-02 |
| organic substance metabolic process (GO:0071704) | 878 | 79 | 55.01 | + | 1.44 | 3.47E-04 | 4.52E-02 |
| biological_process (GO:0008150) | 1496 | 131 | 93.73 | + | 1.4 | 5.45E-08 | 8.54E-05 |
| Unclassified (UNCLASSIFIED) | 1393 | 50 | 87.27 | - | 0.57 | 5.45E-08 | 4.27E-05 |
| GO molecular function complete | Staphylococcus aureus - REFLIST (2889) | p003p (181) | p003p (expected) | p003p (over/under) | p003p (fold Enrichment) | p003p (raw P-value) | p003p (FDR) |
| nucleotide binding (GO:0000166) | 382 | 43 | 23.93 | + | 1.8 | 2.13E-04 | 3.10E-02 |
| nucleoside phosphate binding (GO:1901265) | 382 | 43 | 23.93 | + | 1.8 | 2.13E-04 | 2.75E-02 |
| heterocyclic compound binding (GO:1901363) | 710 | 74 | 44.48 | + | 1.66 | 3.81E-06 | 8.89E-04 |
| organic cyclic compound binding (GO:0097159) | 710 | 74 | 44.48 | + | 1.66 | 3.81E-06 | 7.41E-04 |
| ion binding (GO:0043167) | 569 | 58 | 35.65 | + | 1.63 | 1.33E-04 | 2.22E-02 |
| binding (GO:0005488) | 898 | 88 | 56.26 | + | 1.56 | 2.31E-06 | 6.75E-04 |
| catalytic activity (GO:0003824) | 1105 | 103 | 69.23 | + | 1.49 | 9.48E-07 | 3.68E-04 |
| molecular_function (GO:0003674) | 1567 | 137 | 98.17 | + | 1.4 | 7.99E-09 | 9.31E-06 |
| Unclassified (UNCLASSIFIED) | 1322 | 44 | 82.83 | - | 0.53 | 7.99E-09 | 4.66E-06 |
| PANTHER GO-Slim Molecular Function | Staphylococcus aureus - REFLIST (2889) | p003p (181) | p003p (expected) | p003p (over/under) | p003p (fold Enrichment) | p003p (raw P-value) | p003p (FDR) |
| catalytic activity (GO:0003824) | 498 | 52 | 31.2 | + | 1.67 | 2.01E-04 | 1.65E-02 |
| molecular_function (GO:0003674) | 729 | 72 | 45.67 | + | 1.58 | 3.56E-05 | 4.38E-03 |
| Unclassified (UNCLASSIFIED) | 2160 | 109 | 135.33 | - | 0.81 | 3.56E-05 | 8.76E-03 |
| PANTHER GO-Slim Biological Process | Staphylococcus aureus - REFLIST (2889) | p003p (181) | p003p (expected) | p003p (over/under) | p003p (fold Enrichment) | p003p (raw P-value) | p003p (FDR) |
| nitrogen compound metabolic process (GO:0006807) | 371 | 42 | 23.24 | + | 1.81 | 2.74E-04 | 4.08E-02 |
| cellular metabolic process (GO:0044237) | 425 | 48 | 26.63 | + | 1.8 | 7.05E-05 | 4.20E-02 |
| organic substance metabolic process (GO:0071704) | 415 | 45 | 26 | + | 1.73 | 3.43E-04 | 4.09E-02 |
| metabolic process (GO:0008152) | 443 | 48 | 27.75 | + | 1.73 | 2.16E-04 | 4.30E-02 |
| cellular process (GO:0009987) | 485 | 52 | 30.39 | + | 1.71 | 1.10E-04 | 3.29E-02 |
| PANTHER Protein Class | Staphylococcus aureus - REFLIST (2889) | p003p (181) | p003p (expected) | p003p (over/under) | p003p (fold Enrichment) | p003p (raw P-value) | p003p (FDR) |
| protein class (PC00000) | 986 | 95 | 61.77 | + | 1.54 | 1.16E-06 | 5.99E-05 |
| Unclassified (UNCLASSIFIED) | 1903 | 86 | 119.23 | - | 0.72 | 1.16E-06 | 1.20E-04 |
| GO biological process complete | Staphylococcus aureus - REFLIST (2889) | pyo (146) | pyo (expected) | pyo (over/under) | pyo (fold Enrichment) | pyo (raw P-value) | pyo (FDR) |
| cellular amino acid metabolic process (GO:0006520) | 136 | 19 | 6.87 | + | 2.76 | 1.11E-04 | 1.33E-02 |
| small molecule metabolic process (GO:0044281) | 375 | 45 | 18.95 | + | 2.37 | 4.79E-08 | 9.37E-06 |
| carboxylic acid metabolic process (GO:0019752) | 198 | 23 | 10.01 | + | 2.3 | 4.17E-04 | 3.63E-02 |
| oxoacid metabolic process (GO:0043436) | 202 | 23 | 10.21 | + | 2.25 | 4.69E-04 | 3.67E-02 |
| organic acid metabolic process (GO:0006082) | 222 | 25 | 11.22 | + | 2.23 | 2.56E-04 | 2.50E-02 |
| cellular biosynthetic process (GO:0044249) | 469 | 48 | 23.7 | + | 2.03 | 1.45E-06 | 2.52E-04 |
| organic substance biosynthetic process (GO:1901576) | 476 | 48 | 24.06 | + | 2 | 2.65E-06 | 4.14E-04 |
| biosynthetic process (GO:0009058) | 495 | 49 | 25.02 | + | 1.96 | 3.75E-06 | 5.34E-04 |
| Nucleobase-  containing compound metabolic process (GO:0006139) | 331 | 32 | 16.73 | + | 1.91 | 5.50E-04 | 4.10E-02 |
| organic substance metabolic process (GO:0071704) | 878 | 84 | 44.37 | + | 1.89 | 5.65E-11 | 4.42E-08 |
| cellular metabolic process (GO:0044237) | 837 | 80 | 42.3 | + | 1.89 | 2.49E-10 | 6.50E-08 |
| cellular aromatic compound metabolic process (GO:0006725) | 394 | 37 | 19.91 | + | 1.86 | 2.30E-04 | 2.40E-02 |
| primary metabolic process (GO:0044238) | 748 | 70 | 37.8 | + | 1.85 | 2.87E-08 | 6.41E-06 |
| metabolic process (GO:0008152) | 1030 | 95 | 52.05 | + | 1.83 | 3.57E-12 | 5.59E-09 |
| cellular process (GO:0009987) | 947 | 87 | 47.86 | + | 1.82 | 1.39E-10 | 4.35E-08 |
| heterocycle metabolic process (GO:0046483) | 408 | 37 | 20.62 | + | 1.79 | 4.54E-04 | 3.74E-02 |
| cellular nitrogen compound metabolic process (GO:0034641) | 491 | 43 | 24.81 | + | 1.73 | 3.12E-04 | 2.87E-02 |
| organonitrogen compound metabolic process (GO:1901564) | 514 | 45 | 25.98 | + | 1.73 | 1.79E-04 | 2.00E-02 |
| nitrogen compound metabolic process (GO:0006807) | 717 | 62 | 36.23 | + | 1.71 | 6.12E-06 | 7.98E-04 |
| biological_process (GO:0008150) | 1496 | 115 | 75.6 | + | 1.52 | 7.10E-11 | 2.78E-08 |
| Unclassified (UNCLASSIFIED) | 1393 | 31 | 70.4 | - | 0.44 | 7.10E-11 | 3.71E-08 |
| GO molecular function complete | Staphylococcus aureus - REFLIST (2889) | pyo (146) | pyo (expected) | pyo (over/under) | pyo (fold Enrichment) | pyo (raw P-value) | pyo (FDR) |
| protein dimerization activity (GO:0046983) | 11 | 5 | 0.56 | + | 8.99 | 6.83E-04 | 3.46E-02 |
| protein binding (GO:0005515) | 28 | 9 | 1.42 | + | 6.36 | 4.22E-05 | 3.08E-03 |
| oxidoreductase activity, acting on CH-OH group of donors (GO:0016614) | 41 | 11 | 2.07 | + | 5.31 | 2.41E-05 | 1.87E-03 |
| magnesium ion binding (GO:0000287) | 53 | 11 | 2.68 | + | 4.11 | 1.80E-04 | 1.10E-02 |
| transition metal ion binding (GO:0046914) | 83 | 16 | 4.19 | + | 3.81 | 1.30E-05 | 1.38E-03 |
| coenzyme binding (GO:0050662) | 111 | 16 | 5.61 | + | 2.85 | 2.92E-04 | 1.62E-02 |
| cofactor binding (GO:0048037) | 154 | 22 | 7.78 | + | 2.83 | 2.19E-05 | 2.13E-03 |
| metal ion binding (GO:0046872) | 271 | 37 | 13.7 | + | 2.7 | 5.05E-08 | 9.82E-06 |
| cation binding (GO:0043169) | 276 | 37 | 13.95 | + | 2.65 | 7.83E-08 | 1.30E-05 |
| oxidoreductase activity (GO:0016491) | 182 | 22 | 9.2 | + | 2.39 | 2.43E-04 | 1.42E-02 |
| transferase activity, transferring phosphorus-  containing groups (GO:0016772) | 161 | 19 | 8.14 | + | 2.34 | 8.91E-04 | 4.33E-02 |
| ion binding (GO:0043167) | 569 | 62 | 28.76 | + | 2.16 | 1.59E-09 | 3.71E-07 |
| small molecule binding (GO:0036094) | 427 | 46 | 21.58 | + | 2.13 | 8.69E-07 | 1.27E-04 |
| nucleotide binding (GO:0000166) | 382 | 39 | 19.3 | + | 2.02 | 2.30E-05 | 2.06E-03 |
| nucleoside phosphate binding (GO:1901265) | 382 | 39 | 19.3 | + | 2.02 | 2.30E-05 | 1.91E-03 |
| transferase activity (GO:0016740) | 366 | 37 | 18.5 | + | 2 | 5.00E-05 | 3.43E-03 |
| drug binding (GO:0008144) | 311 | 31 | 15.72 | + | 1.97 | 3.83E-04 | 2.03E-02 |
| anion binding (GO:0043168) | 378 | 37 | 19.1 | + | 1.94 | 1.03E-04 | 6.69E-03 |
| binding (GO:0005488) | 898 | 85 | 45.38 | + | 1.87 | 7.44E-11 | 2.17E-08 |
| catalytic activity (GO:0003824) | 1105 | 101 | 55.84 | + | 1.81 | 2.43E-13 | 9.43E-11 |
| heterocyclic compound binding (GO:1901363) | 710 | 62 | 35.88 | + | 1.73 | 5.52E-06 | 7.15E-04 |
| organic cyclic compound binding (GO:0097159) | 710 | 62 | 35.88 | + | 1.73 | 5.52E-06 | 6.43E-04 |
| molecular_function (GO:0003674) | 1567 | 128 | 79.19 | + | 1.62 | 3.24E-17 | 3.78E-14 |
| Unclassified (UNCLASSIFIED) | 1322 | 18 | 66.81 | - | 0.27 | 3.24E-17 | 1.89E-14 |
| GO cellular component complete | Staphylococcus aureus - REFLIST (2889) | pyo (146) | pyo (expected) | pyo (over/under) | pyo (fold Enrichment) | pyo (raw P-value) | pyo (FDR) |
| cytoplasm (GO:0005737) | 332 | 39 | 16.78 | + | 2.32 | 8.66E-07 | 1.01E-04 |
| intracellular (GO:0005622) | 411 | 42 | 20.77 | + | 2.02 | 1.21E-05 | 7.08E-04 |
| PANTHER Pathways | Staphylococcus aureus - REFLIST (2889) | pyo (146) | pyo (expected) | pyo (over/under) | pyo (fold Enrichment) | pyo (raw P-value) | pyo (FDR) |
| Unclassified (UNCLASSIFIED) | 2663 | 117 | 134.58 | - | 0.87 | 8.09E-06 | 5.67E-04 |
| PANTHER GO-Slim Molecular Function | Staphylococcus aureus - REFLIST (2889) | pyo (146) | pyo (expected) | pyo (over/under) | pyo (fold Enrichment) | pyo (raw P-value) | pyo (FDR) |
| catalytic activity (GO:0003824) | 498 | 53 | 25.17 | + | 2.11 | 1.10E-07 | 2.72E-05 |
| molecular_function (GO:0003674) | 729 | 66 | 36.84 | + | 1.79 | 3.66E-07 | 4.50E-05 |
| Unclassified (UNCLASSIFIED) | 2160 | 80 | 109.16 | - | 0.73 | 3.66E-07 | 3.00E-05 |
| PANTHER GO-Slim Biological Process | Staphylococcus aureus - REFLIST (2889) | pyo (146) | pyo (expected) | pyo (over/under) | pyo (fold Enrichment) | pyo (raw P-value) | pyo (FDR) |
| metabolic process (GO:0008152) | 443 | 44 | 22.39 | + | 1.97 | 1.31E-05 | 7.82E-03 |
| cellular metabolic process (GO:0044237) | 425 | 40 | 21.48 | + | 1.86 | 1.30E-04 | 1.55E-02 |
| organic substance metabolic process (GO:0071704) | 415 | 39 | 20.97 | + | 1.86 | 1.82E-04 | 1.80E-02 |
| cellular process (GO:0009987) | 485 | 45 | 24.51 | + | 1.84 | 4.68E-05 | 1.40E-02 |
| biological_process (GO:0008150) | 606 | 52 | 30.63 | + | 1.7 | 7.70E-05 | 1.53E-02 |
| Unclassified (UNCLASSIFIED) | 2283 | 94 | 115.37 | - | 0.81 | 7.70E-05 | 1.15E-02 |
| PANTHER GO-Slim Cellular Component | Staphylococcus aureus - REFLIST (2889) | pyo (146) | pyo (expected) | pyo (over/under) | pyo (fold Enrichment) | pyo (raw P-value) | pyo (FDR) |
| cytoplasm (GO:0005737) | 265 | 30 | 13.39 | + | 2.24 | 4.36E-05 | 4.05E-03 |
| cytoplasmic part (GO:0044444) | 184 | 20 | 9.3 | + | 2.15 | 1.82E-03 | 4.22E-02 |
| intracellular part (GO:0044424) | 288 | 31 | 14.55 | + | 2.13 | 8.13E-05 | 3.78E-03 |
| intracellular (GO:0005622) | 292 | 31 | 14.76 | + | 2.1 | 1.46E-04 | 4.54E-03 |
| PANTHER Protein Class | Staphylococcus aureus - REFLIST (2889) | pyo (146) | pyo (expected) | pyo (over/under) | pyo (fold Enrichment) | pyo (raw P-value) | pyo (FDR) |
| metabolite interconversion enzyme (PC00262) | 473 | 50 | 23.9 | + | 2.09 | 4.06E-07 | 1.39E-05 |
| protein class (PC00000) | 986 | 82 | 49.83 | + | 1.65 | 1.68E-07 | 1.73E-05 |
| Unclassified (UNCLASSIFIED) | 1903 | 64 | 96.17 | - | 0.67 | 1.68E-07 | 8.64E-06 |

References (supplemental)

1.	Lees JA, Vehkala M, Välimäki N, Harris SR, Chewapreecha C, Croucher NJ, Marttinen P, Davies MR, Steer AC, Tong SYC, Honkela A, Parkhill J, Bentley SD, Corander J. 2016. Sequence element enrichment analysis to determine the genetic basis of bacterial phenotypes. Nat Commun 7.
